# Lysosomal activation in bladder epithelium enhances intracellular antibiotic clearance of uropathogenic *Escherichia coli*

**DOI:** 10.1101/2025.10.22.683857

**Authors:** Kathrin Tomasek, Kristina Skurvydaite, Gauri Paduthol, Allison M. Burns, Léa Schlunke, Valentin Borgeat, Christian Pasquali, Mario Romani, John D. McKinney

**Affiliations:** Laboratory of Microbiology and Microtechnology, School of Life Sciences (SV), École Polytechnique Fédérale de Lausanne, 1015 Lausanne, Switzerland; Bioinformatics Competence Centre, École Polytechnique Fédérale de Lausanne, 1015 Lausanne, Switzerland; OM Pharma Ltd., Rue du Bois-du-Lan 22, 1217 Meyrin / Geneva, Switzerland

## Abstract

Recurrent urinary tract infections (UTIs) are a major clinical burden, driven in part by the ability of uropathogenic Escherichia coli (UPEC) to establish intracellular niches within the bladder epithelium, where bacteria withstand antibiotics and host defenses. The oral bacterial lysate OM-89 (Uro-Vaxom®), a clinically approved and globally used therapy for the prevention and management of recurrent UTIs for several decades, reduces recurrence rates, but its cellular mechanisms of action remain incompletely understood. Here, we demonstrate that OM-89 strengthens antimicrobial defenses in bladder epithelial cells and, in combination with antibiotic therapy, limits post-treatment regrowth in epithelial infection models. Using bladder organoids together with differentiated epithelial monolayers, OM-89 promotes lysosomal acidification and increases lysosomal protease activity, driving intracellular UPEC toward degradative compartments. In parallel, OM-89 improves intracellular antibiotic efficacy across multiple antibiotic classes, leading to enhanced bacterial clearance and reduced post-treatment bacterial recovery. These effects are conserved across distinct UPEC strains and in both murine and human epithelial models. Our findings position the bladder epithelium from a passive barrier to an active, targetable determinant of treatment outcome and suggest host-directed modulation of epithelial antimicrobial pathways as a promising strategy to enhance intracellular bacterial clearance.

## Introduction

Intracellular persistence of bacterial pathogens represents a major barrier to successful infection clearance, as antibiotics often fail to efficiently accumulate within host cells (Chifiriuc et al, 2016; Kamaruzzaman et al, 2017) and due to the pathogens’ ability to escape degradative compartments (Naskar et al, 2023; Miao et al, 2015). This limitation is particularly relevant for mucosal epithelia, which act as the first physical barrier during many infections. Whether epithelial cell-intrinsic antimicrobial pathways can be therapeutically reinforced to enhance intracellular bacterial clearance remains insufficiently explored.

Urinary tract infections (UTIs) exemplify this challenge (Medina & Castillo-Pino, 2019; Stracy et al, 2022; Foxman et al, 2000). Uropathogenic *Escherichia coli* (UPEC), the predominant pathogen of UTIs, invades bladder epithelial cells and establishes protected intracellular niches that promote persistence and recurrence (Mysorekar & Hultgren, 2006). These include rapidly replicating intracellular bacterial communities (Anderson et al, 2004) and long-lived quiescent reservoirs (Mulvey et al, 2001), both of which are shielded from immune responses and incompletely eradicated by antibiotics (Mysorekar & Hultgren, 2006). Recurrent UTIs therefore drive repeated antibiotic exposure, accelerating antimicrobial resistance and increasing the risk of treatment failure (Murray et al, 2022). Despite the clear role of intracellular bacterial reservoirs in the recurrence of UTIs, strategies aimed at strengthening epithelial antimicrobial mechanisms rather than directly targeting UPEC remain rare.

The bladder epithelium is an immunologically active tissue capable of pathogen sensing, trafficking control and lysosomal degradation (Abraham & Miao, 2015). For example, bladder epithelial cells expel invading UPEC via fusiform vesicles or traffic invading bacteria via endo-lysosomal pathways or autophagosomes into degradative compartments and secrete antimicrobial peptides into the urine (Hou et al, 2025). Experimental and clinical data suggest that modulating mucosal immunity can reduce recurrent UTIs (Pei et al, 2025). While this observation highlights the coordinated contribution of epithelial barriers and mucosal immune cells to host defense, the epithelial cell-intrinsic effector mechanisms underlying protection remain poorly defined.

OM-89 (Active Pharmaceutical Ingredient of Uro-Vaxom®), an orally administered lysate derived from 18 different *E. coli* strains, is clinically approved and has been used for nearly four decades for the prevention of recurrent UTIs (Frey et al, 1986; Bauer et al, 2005). OM-89 is approved in numerous countries and is widely prescribed as prophylaxis in patients with recurrent lower UTIs, either alone or in combination with antibiotic therapy. Clinical trials have demonstrated that OM-89 significantly reduces recurrence rates (Magasi et al, 1994; Schulman et al, 1993; Volontè et al, 2025) and experimental studies suggest that it exerts immunomodulatory effects through activation of innate immune cells and stimulation of antimicrobial humoral responses (Rosenthal, 1986; Huber et al, 2000; Schmidhammer et al, 2002). However, despite extensive clinical use and experimental investigation, the cellular mechanisms underlying its protective effects remain incompletely understood. For instant, rodent infection studies have demonstrated protective effects of OM-89 (Bosch et al, 1988; Lee et al, 2006), including when combined antibiotic therapy (Canton et al, 2025; Bessler et al, 2010) but this observed *in vivo* protection could not be linked to any major quantitative changes in bladder immune cell infiltration (Canton et al, 2025), leaving the underlying molecular mechanism unresolved. However, Canton et al. speculated that the bladder epithelium itself may act as a direct effector site, supported by pharmacokinetic evidence showing urinary accumulation of OM-89-derived components (van Dijk, 1982). In parallel, other bacterial lysates have been shown to directly modulate epithelial responses in the respiratory tract (Roth et al, 2017; Sidoti Migliore et al, 2023; Pivniouk et al, 2022), supporting the concept of epithelial conditioning across different mucosal surfaces. Together, these observations raise the possibility that the bladder epithelium, and specifically epithelial cell-intrinsic antimicrobial pathways, may contribute to OM-89-mediated protection.

Here, we investigated whether OM-89 directly modulates antimicrobial pathways in bladder epithelial cells using a previously established murine bladder organoid model (Sharma et al, 2021), a human-derived bladder organoid model and differentiated epithelial monolayers that recapitulate urothelial differentiation. We show that OM-89 remodels endo-lysosomal networks and enhances lysosomal acidification and protease activity, indicating that epithelial antimicrobial pathways can be pharmacologically reinforced to enhance intracellular bacterial clearance. Together with increased intracellular accumulation of antibiotics across different classes, this leads to improved intracellular killing and reduced bacterial regrowth across diverse UPEC strains. The key features of this phenotype, including lysosomal expansion, acidification, protease activation and enhanced antibiotic uptake, are conserved in human bladder epithelial monolayers and organoid cultures, despite species-specific differences in autophagic flux regulation. Together, these findings reveal a previously unrecognized epithelial lysosome-centered mechanism by which OM-89 enhances intracellular antibiotic performance and repositions the bladder epithelium from a passive reservoir of infection reactivation to an actively transformable antimicrobial compartment influencing treatment outcomes.

## Results

### OM-89 enhances epithelial control of intracellular UPEC and limits post-treatment bacterial regrowth

To determine whether OM-89 enhances bladder epithelial control of intracellular UPEC during antibiotic treatment, we used a previously established and characterized mouse bladder organoid model (Sharma et al, 2021) infected with the well-characterized UPEC strain CFT073 (Mobley et al, 1990), originally isolated from a pyelonephritis patient. We monitored bacterial growth, antibiotic-mediated killing and post-treatment regrowth within the organoids. To model clinically relevant treatment scenarios with the clinically used therapy Uro-Vaxom®, we applied OM-89 under three different regimens (Figure 1A): (i) pre-application – OM-89 exposure for 72 hours followed by a 24-hour rest period before microinjection of UPEC; (ii) co-application – OM-89 exposure initiated at the same time as antibiotic treatment; and (iii) continuous application – OM-89 exposure both 72 hours before microinjection and throughout the entire experiment. We quantified UPEC fluorescence area overlapping with the organoid area as a proxy for viable burden and regrowth, normalized within experiments to PBS controls. Notably, OM-89 did not influence bacterial loads during the initial infection phase compared to the PBS control groups (4h post-infection (pi); SI Figure 1A-C), indicating no antimicrobial effects of OM-89 alone. However, OM-89 significantly reduced bacterial regrowth following post-antibiotic treatment (10h pi; 10x MIC of ampicillin) compared to the PBS control in all three treatment regimens (Figure 1B-D), with the effect being most pronounced when OM-89 was present during the antibiotic treatment phase (co-application, Figure 1C; and continuous application, Figure 1D). These positive effects persisted up to eight hours post-antibiotic withdrawal (15h pi) in the continuous application regimen (Figure 1E).

**Figure 1.**
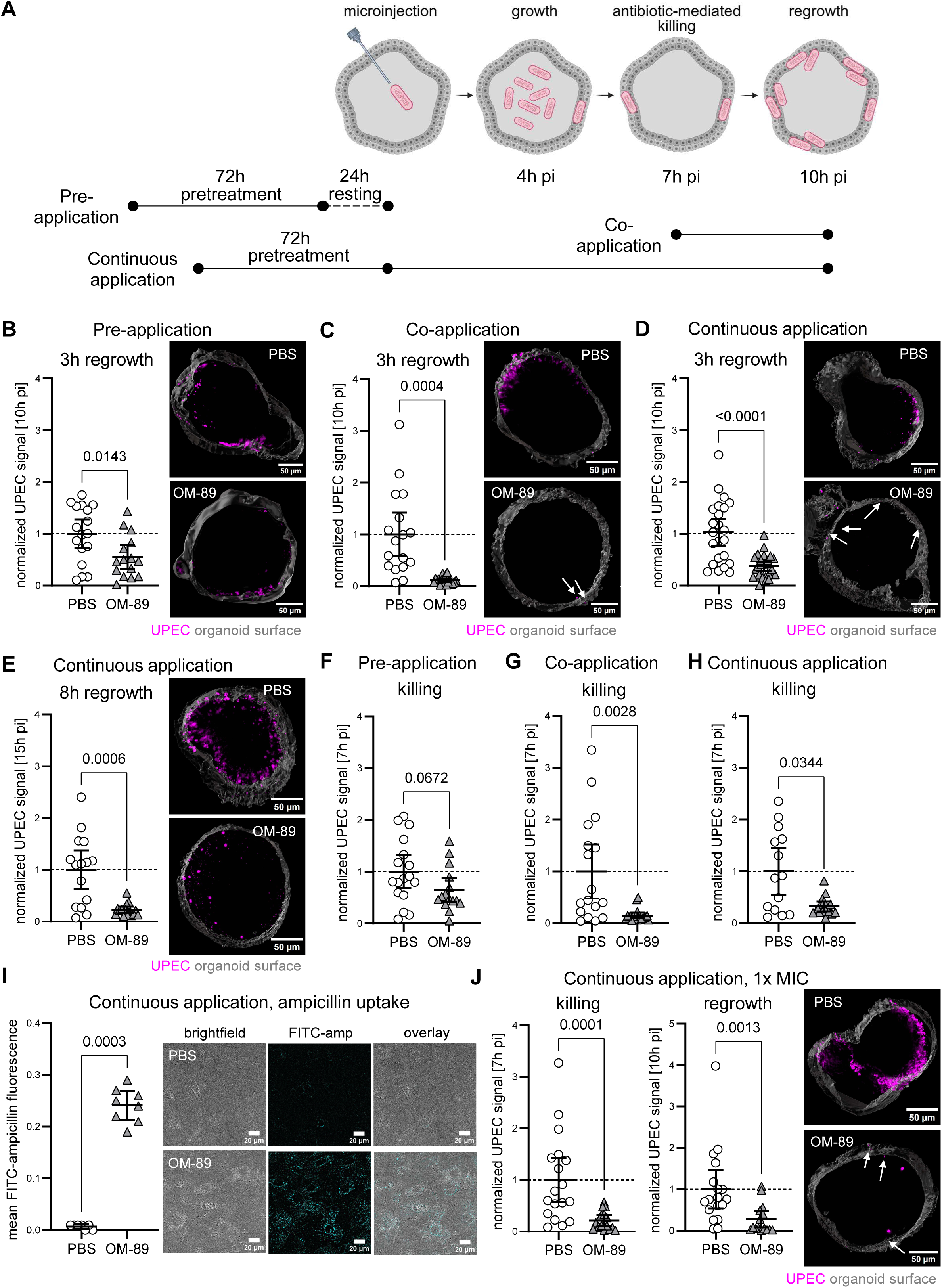
OM-89 reduces regrowth of CFT073 in mouse bladder organoids due to enhanced antibiotic-mediated killing. (A) Mouse bladder organoids treated with OM-89 under different regimes (pre-application: OM-89 exposure for 72h and 24h rest period; co-application: OM-89 exposure together with antibiotic treatment; continuous application: OM-89 exposure both 72h before microinjection and throughout the entire experiment). After microinjection of fluorescently labelled UPEC (0h post-infection, pi), bacterial growth (4h pi), antibiotic-mediated killing (7h pi), and regrowth 3 hours after antibiotic withdrawal (10h pi) were monitored. Created with BioRender.com. (B)-(D) CFT073 signal 10h pi (3h regrowth) at 10x MIC in (B) pre-application, (C) coapplication, and (D) continuous application regimes in mouse bladder organoids. Each dot represents one organoid. Values normalized to PBS control (dashed line). Mean ± 95% CI. Welch’s t test. N ≥ 15 per condition for (B), n ≥ 17 per condition for (C) and n ≥ 23 per condition for (D). Images show representative organoids (grey, surface reconstruction) and UPEC (magenta, some marked with arrows). (E) CFT073 signal 15h pi (8h regrowth) at 10x MIC in the continuous application regime in mouse bladder organoids. Values normalized to PBS control (dashed line). Mean ± 95% CI. Mann-Whitney test. N ≥ 15 per condition. Images show representative organoids (grey, surface reconstruction) and UPEC (magenta, some marked with arrows). (F)-(H) CFT073 signal 7h pi (killing) at 10x MIC in (F) pre-application, (G) co-application, and (H) continuous application regimes in mouse bladder organoids. Values normalized to PBS control (dashed line). Mean ± 95% CI. Welch’s t test for (F), Mann-Whitney test for (G), (H). N ≥ 15 per condition for (F), n ≥ 17 per condition for (G) and n ≥ 14 per condition for (H). (I) Uptake of FITC-labelled ampicillin into monolayers of mouse bladder epithelial cells during the continuous application regime. Quantification of fluorescence after background subtraction of PBS- or OM-89-treated cells without labelled antibiotic. Each dot represents one field of view. Mean ± 95% CI. Mann-Whitney test. N ≥ 7 per condition. Z-projection (maximum intensity) of representative images. FITC-ampicillin shown in cyan. (J) CFT073 signal 7h pi (1x MIC; killing and 10h pi (regrowth) in mouse bladder organoids during the continuous application regime. Each dot represents one organoid. Values normalized to PBS control (dashed line). Mean ± 95% CI. Mann-Whitney test. N ≥ 15 per condition. Images show representative organoids (grey, surface reconstruction) and UPEC (magenta, some marked with arrows).

We next examined the effect of OM-89 during antibiotic treatment (7h pi). While the pre-application regimen showed modest modulation of antibiotic effects (Figure 1F), bacterial loads within organoids were significantly reduced in conditions where OM-89 was present at the time of antibiotic treatment (co-application, Figure 1G; continuous application, Figure 1H). Following bacterial burden in individual organoids in the continuous application regimen over time, we observed that OM-89 consistently enhanced bacterial clearance throughout antibiotic treatment and regrowth phases (SI Figure 1D). Finally, as observed previously (Sharma et al, 2021), we found that regrowth of UPEC occurred preferentially from the organoid wall – irrespective of treatment with OM-89 (SI Figure 1E, SI movie 1 and 2).

To further investigate OM-89’s modulating effect, we focused on the continuous application regimen for the rest of the manuscript as this treatment regime most likely reflects the clinical application in patients. First, given the enhanced antibiotic-mediated killing in OM-89-treated cells, we asked if OM-89 could alter antibiotic accumulation inside bladder epithelial cells, as many conventional antibiotics, including β-lactam antibiotics such as ampicillin, display low intracellular accumulation levels (Carryn et al, 2003). We therefore examined the uptake of fluorescently labeled ampicillin into differentiated monolayers of mouse bladder epithelial cells (SI Figure 2) and observed significantly increased accumulation of fluorescently labelled ampicillin in bladder epithelial cells treated with OM-89 (Figure 1I). Importantly, UPEC infection alone did not increase intracellular antibiotic accumulation compared to uninfected controls (SI Figure 1F), indicating that enhanced uptake is a specific response to OM-89 exposure rather than a general *E. coli*-driven stimulation response. Furthermore, the increased antibiotic uptake was predominantly observed in umbrella-like cells (cytokeratin (CK)20 positive cells), whereas intermediate cells (CK13 positive cells) showed little uptake (SI Figure 1G), identifying terminally differentiated cells as the main contributors to the uptake phenotype. The effect of increased antibiotic uptake extended to gentamicin, an aminoglycoside antibiotic with usually slow and insufficient intracellular accumulation dynamics commonly used in infection assays to eradicate only extracellular bacteria (Kadurugamuwa & Beveridge, 1998; Carryn et al, 2003; Elsinghorst, 1994) (SI Figure 1H). It is important to note that the TFP ester of the fluorescent dye used to label ampicillin and gentamicin preferentially reacts with the non-protonated form of free amine groups, which may alter the antibiotics’ intracellular behavior and potentially affect their antimicrobial activity. Nonetheless, OM-89 treatment led to increased intracellular accumulation of both fluorescently labelled ampicillin and gentamicin, as well as Dextran-TMR, a polysaccharide uptake marker (SI Figure 1I), suggesting a generalized enhancement of cellular uptake in bladder epithelial cells compared to PBS-treated controls. However, because the FITC labeling may alter antibiotic behavior, our uptake measurements reflect fluorescent proxies rather than native drug activity.

**Figure 2.**
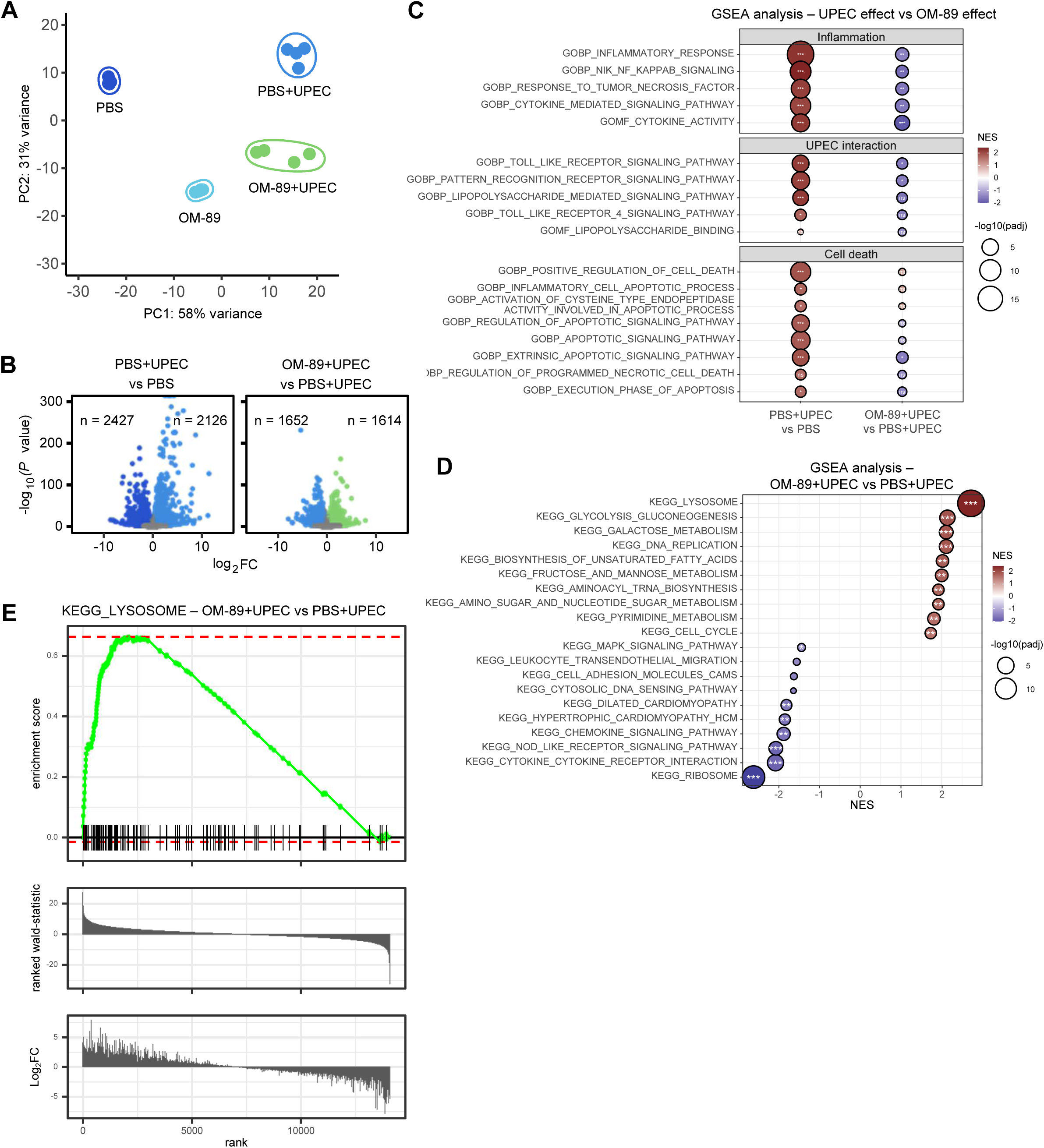
OM-89 modulates inflammatory signaling and promotes lysosomal pathways in bladder epithelial cells. (A) Principal component analysis (PCA) of mouse bladder organoids treated with OM-89 or PBS, with and without CFT073 infection (PBS – dark blue, OM-89 – cyan, PBS+UPEC – light blue, OM-89+UPEC – green). (B) Volcano plots of differentially expressed genes (log_2_FC ≥ 1; adjusted p ≤ 0.05) comparing PBS-treated organoids with (light blue) and without infection (dark blue), and OM-89- (green) versus PBS-treated organoids during infection (light blue). (C) Gene set enrichment analysis (GSEA) of treatment and infection groups. GO-BP terms were clustered into infection-relevant pathways. NES, normalized enrichment score. Nonsignificant changes not highlighted. (D) GSEA of the top 10 Kyoto Encyclopedia of Genes and Genomes (KEGG) terms comparing OM-89- and PBS-treated organoids during infection. Ns changes not highlighted (E) Enrichment plot of lysosomal pathway genes (KEGG term) in OM-89- versus PBS-treated organoids during infection (FDR corrected p-value = 1.88e-14 and uncorrected pvalue = 1.21e-16). (A)-(E) Data from four independent wells with n ≥ 50 organoids each.

To determine whether the increased uptake of antibiotics contributed to OM-89’s protective effect, we reduced the antibiotic concentration from 10-fold to 1-fold the MIC. Even under these conditions, OM-89 maintained its ability to significantly enhance bacterial killing within three hours (7h pi) and suppress regrowth up to three hours post-treatment (10h pi) (Figure 1J), suggesting that OM-89 improves antibiotic efficacy potentially by increasing intracellular accumulation.

Together, these data show that OM-89 potentiates antibiotic activity within bladder epithelial cells, enabling improved bacterial clearance within bladder organoids and sustained suppression of post-treatment bacterial regrowth, consistent with enhanced intracellular activity.

### OM-89 reroutes epithelial degradative pathways toward a lysosome-dominant antimicrobial state

To define how OM-89 alters epithelial antimicrobial pathways during infection, we performed transcriptomic profiling of infected and treated bladder organoids.

Principal component analysis revealed a strong separation between control and infected organoids (Figure 2A), consistent with widespread differential gene expression across conditions (Figure 2B, SI Figure 3A). Gene set enrichment analysis (GSEA) confirmed that CFT073 infection induced pathways associated with inflammation, bacterial immune signaling, and cell death, whereas OM-89 treatment attenuated the magnitude of these infection-associated signatures (Figure 2C, SI Figure 3B). Notably, OM-89-treated organoids clustered closer to infected samples than to PBS controls (Figure 2A), indicating that OM-89 induces a transcriptional program partially overlapping with infection-responsive pathways. This overlap is further supported by elevated cellular responses to pathogenic stimuli in OM-89-treated organoids compared to uninfected controls (SI Figure 3C). However, OM-89-treated organoids following infection remained transcriptionally distinct from PBS-infected samples, suggesting that OM-89 does not simply recapitulate an infection state but reconfigures epithelial responses in a qualitatively distinct, host defense-oriented manner.

**Figure 3.**
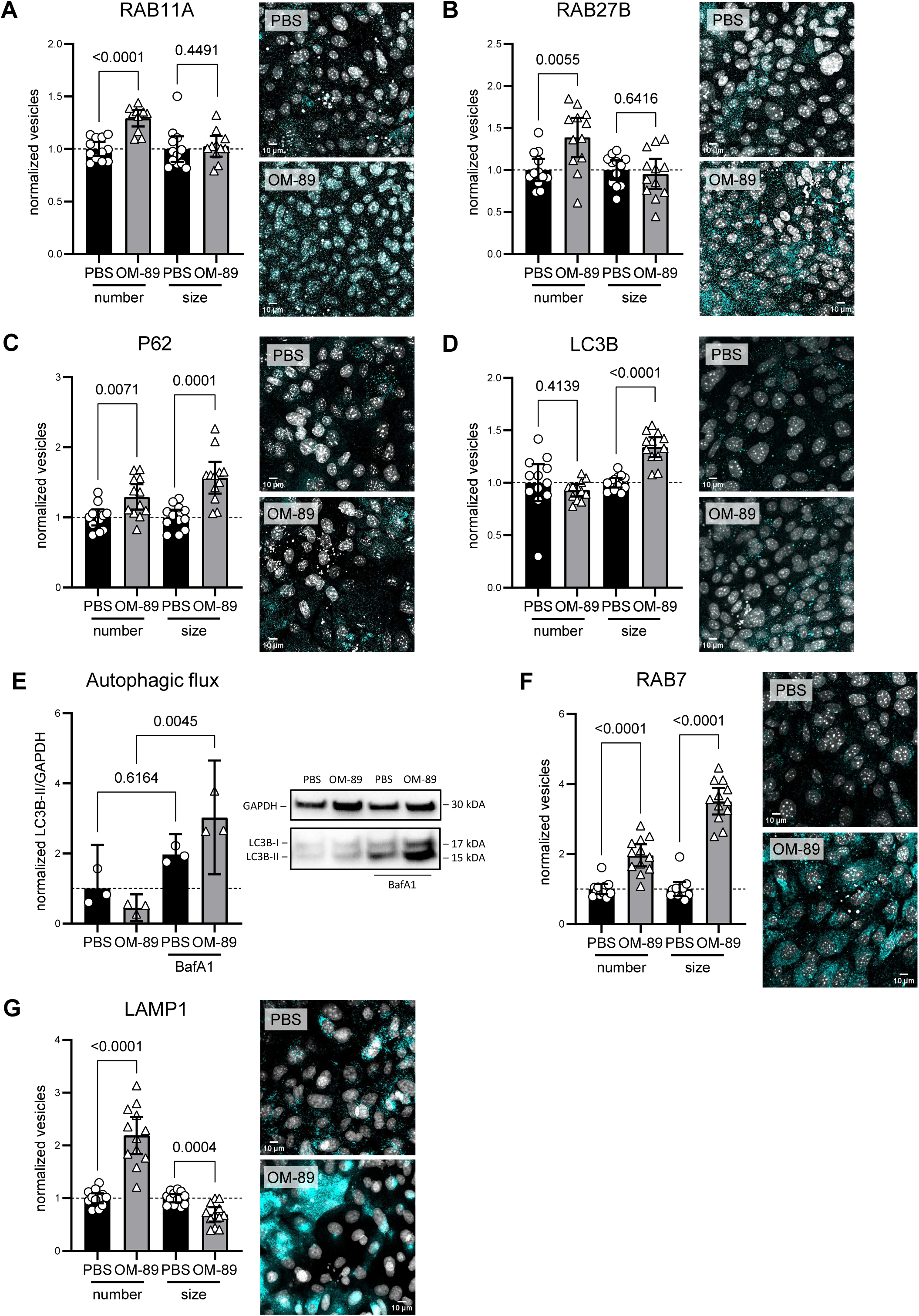
OM-89 remodels vesicle organization and turnover in bladder epithelial cells during UPEC infection. (A)-(D), (F), (G) Quantification of vesicle numbers and size for (A) RAB11A, (B) RAB27B, (C) P62, (D) LC3B, (F) RAB7, and (G) LAMP1 in infected monolayers of mouse bladder epithelial cells with and without OM-89 treatment. Values normalized to PBS control (dashed line). Mean ± 95% CI. Mann-Whitney test for RAB11A and RAB7 (A), (F); Welch’s t test for RAB27B, P62, LC3B and LAMP1 (B)-(D), (G). N = 12 per condition. Z-projection (maximum intensity) of representative images. Foci, cyan; DAPI, grey. (E) Western blot analysis of LC3B during infection of monolayers. LC3B-II/GAPDH ratio normalized to PBS control (dashed line). Mean ± 95% CI. Kruskal-Wallis test with Dunn’s correction. Autophagic flux (BafA1-blocked minus unblocked LC3B-II/GAPDH): PBS = 24.75 ± 10.68; OM-89 = 65.87 ± 9.11. N = 3 per condition

GSEA analysis additionally indicated that the lysosomal pathway was the most significantly induced gene signature following OM-89 treatment during infection (Figure 2D) and further gene set enrichment analysis of lysosomal genes confirmed their involvement in OM-89’s response to infection (Figure 2E). Although, lysosomal pathways were also among the top 10 upregulated gene sets in uninfected organoids exposed to OM-89 (SI Figure 3C), the strong upregulation of lysosome-associated terms, together with autophagy-related pathways, was driven by the combination of infection and OM-89 treatment, as infection in the PBS control group did not upregulate these pathways (SI Figure 3B). These data suggest that OM-89 promotes pathways linked to intracellular cargo degradation, particularly under infectious conditions.

We next examined whether OM-89 modulates intracellular trafficking of UPEC in differentiated monolayers of mouse bladder epithelial cells during infection. We first focused on the endosomal pathway by analyzing Rab11A and Rab27B, two small GTPases implicated in UPEC expulsion from bladder epithelial cells (Miao et al, 2017). Rab11A marks recycling endosomes (Cox et al, 2000), while Rab27B is involved in endosomal secretion (Izumi, 2007) and late endosome/lysosome trafficking (Underwood et al, 2020). OM-89 treatment significantly increased the number of Rab11A- and Rab27B-positive vesicles, without affecting vesicle size (Figure 3A, 3B).

To assess autophagy responses to OM-89 during UPEC infection, we next quantified canonical autophagy markers (Tanida et al, 2008; Kumar et al, 2022). OM-89 significantly increased both the number and size of ubiquitin-binding protein p62 vesicles (Figure 3C), suggesting enhanced cargo sequestration into autophagosomes. In parallel, OM-89 did not significantly change the number of microtubule-associated protein 1A/1B-light chain 3B (LC3b) vesicles, but it significantly increased LC3b vesicle size (Figure 3D), indicating the formation of larger autophagosomes. To determine whether these morphological changes translated into altered autophagic flux during UPEC infection and OM-89 treatment, we performed immunoblotting for membrane-bound LC3b-II in the presence and absence of V-ATPase inhibitor bafilomycin A1 (BafA1) blocking the autophagosome-lysosome fusion. In PBS-treated bladder epithelial cells, UPEC infection stalled flux, as membrane-bound LC3b-II levels did not increase further upon lysosomal inhibition (Figure 3E), consistent with previously described mechanisms of UPEC-mediated autophagy blockade (Li et al, 2024). In contrast, OM-89 treatment overruled this pathogen-induced stalling, as LC3b-II accumulated significantly with BafA1, demonstrating restoration of autophagic flux upon OM-89 treatment.

Finally, we assessed late endosomal (Vanlandingham & Ceresa, 2009) and lysosomal markers (Eskelinen, 2006). OM-89 treatment substantially increased both the number and size of Rab7-positive vesicles, consistent with the expansion of the late endosomal compartment (Figure 3F). Lysosomal-associated membrane protein 1 (Lamp1)-positive vesicles were also strongly increased in number; however, their size was reduced (Figure 3G), potentially indicating remodeling of the lysosomal network.

Together, these findings show that OM-89 shifts epithelial trafficking toward a degradative state by restoring autophagic flux and expanding lysosome-associated compartments during UPEC infection.

### Lysosomal activation is required for OM-89-mediated control of intracellular bacteria

To test whether lysosomal activation could mechanistically explain OM-89-mediated protection, we first used Genebridge analysis (Li et al., 2019) to examine how the lysosomal gene signature identified in our RNA-seq data relates to host defense programs in the human bladder. Module-Module association analysis performed on eight human bladder datasets indicated that the lysosome module has strong positive associations with immune response and bacterial defenses modules (Figure 4A), highlighting a functional link between lysosomal activity and immune defense pathways in the bladder epithelium.

**Figure 4.**
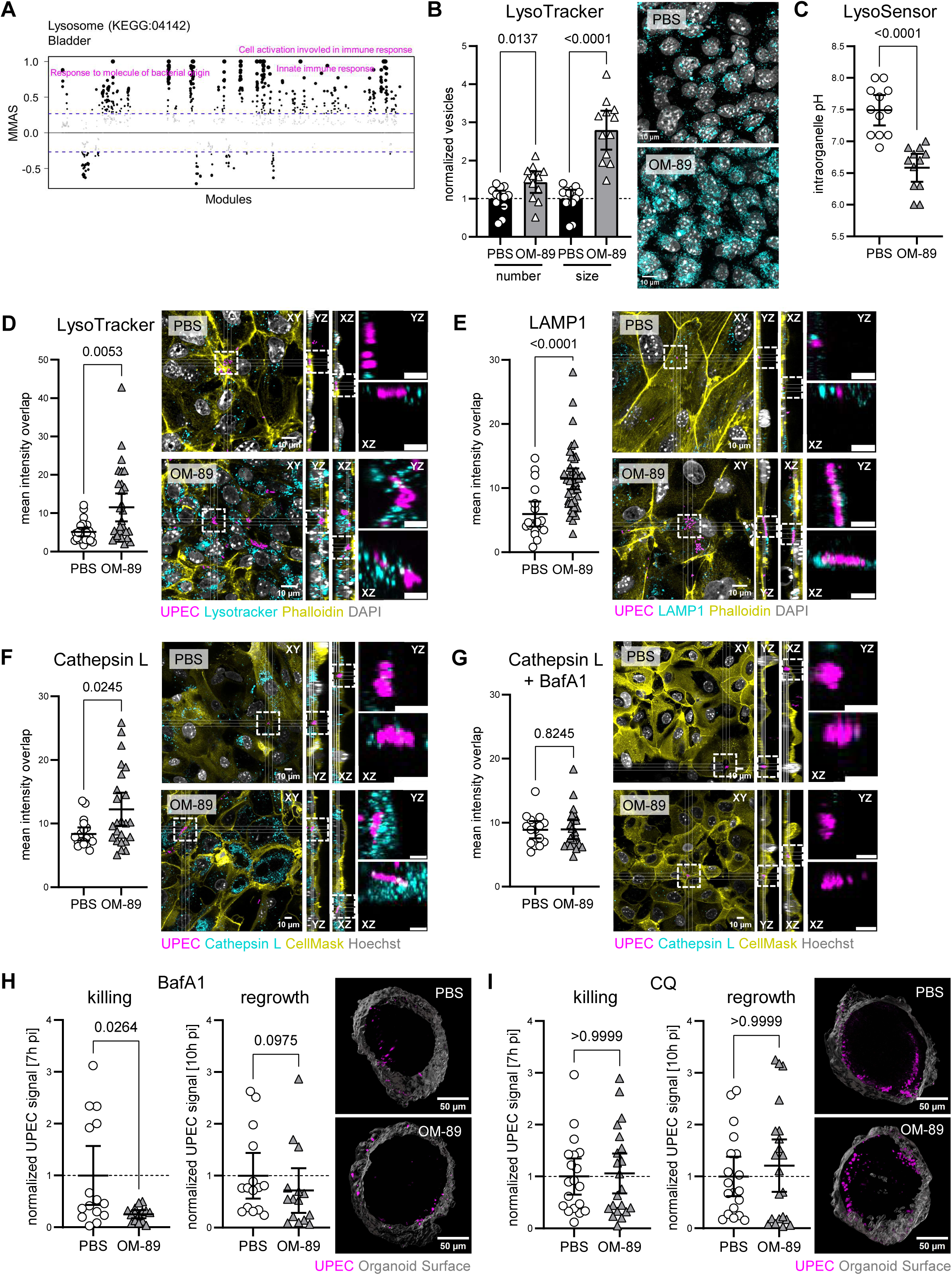
OM-89 enhances lysosomal activity and acidification, while disruption of acidification alters UPEC clearance. (A) GeneBridge MMAS (Module-Module Association Score) analysis of KEGG lysosomal terms in human bladder datasets. Positively correlated modules linked to immune responses and bacterial defense are highlighted. (B) Quantification of vesicle numbers and size of LysoTracker DND-99 in infected monolayers of mouse bladder epithelial cells with and without OM-89 treatment. Values normalized to PBS controls (dashed line). Mean ± 95% CI. Vesicle numbers analyzed with Welch’s t test; vesicle size with Mann-Whitney test. N = 12 per condition. (C) Intraorganelle pH in infected monolayers with and without OM-89 treatment. Mean ± 95% CI. Welch’s t test. N = 12 per condition. (D)-(G) Colocalization of intracellular UPEC with (D) LysoTracker, (E) LAMP1, (F) Cathepsin L and (G) Cathepsin L with bafilomycin A1 (BafA1) blocking in monolayers of mouse bladder epithelial cells. For quantification: Intensities normalized to ROI area drown around intracellular UPEC (see SI Figure 4B-D, F). Mann-Whitney test. N ≥ 24 per condition for LysoTracker, n ≥ 18 per condition for LAMP1, n ≥ 19 per condition for Cathepsin L, n ≥ 15 per condition for Cathepsin L + BafA1 blocking. Enrichment score (= observed overlap/random overlap) between OM-89 vs PBS for (D) 0.68 vs 0.71, (E) 0.94 vs 0.89, (F) 1.25 vs 0.91 and (G) 1.06 vs 1.17. Random overlap from SI Figure 4B-D, F. For images: Cross-section of representative images focused on intracellular bacteria. Intracellularity of bacteria confirmed by inspecting each position visually in XYZ with either phalloidin (D, E) or CellMask (F, G) staining. White inserts in cross-section (XYZ): intracellular UPEC with (D) LysoTracker, (E) LAMP1, (F) Cathepsin L and (G) Cathepsin L with bafilomycin A1 (BafA1) in YZ and XZ. Scale bar of inserts 5 μm. UPEC, magenta; LysoTracker/LAMP1/Cathepsin L, cyan; Phalloidin or CellMask, yellow; DAPI or Hoechst, grey. (H), (I) CFT073 signal 7h pi (killing) and 10 h pi (regrowth) in mouse bladder organoids after blocking with (H) BafA1 or (I) CQ. Each dot represents one organoid. Values normalized to PBS control (dashed line). Mean ± 95% CI. Mann-Whitney test. N = 15 per condition. Images show representative organoids (grey, surface reconstruction) and UPEC (magenta, some marked with arrows).

To further investigate the role of lysosomal modulation in OM-89 mediated effects against UPEC infection, we examined the involvement of lysosomal activity and intracellular pH. In line with remodeling of the late endosomal and lysosomal network (Figure 3F, 3G), OM-89 treatment led to a significant increase in the number of enlarged acidic vesicles (Figure 4B), accompanied by a significant reduction in overall intracellular pH (Figure 4C). Notably, while CFT073 infection elevated intraorganellar pH in PBS-treated controls (pH of 7.5 ± 0.38) compared to uninfected controls (pH of 7.0 ± 0.24), consistently with pathogen-mediated disruption of vesicle acidification (Naskar et al, 2023; Miao et al, 2015), OM-89 treatment maintained a more acidic environment in both conditions (pH of 6.58 ± 0.35 during infection and 6.45 ± 0.28 without infection; Figure 4C, SI Figure 4A). When focusing on intracellular UPEC, OM-89-treated cells showed a significant increase in signal intensity of acidic compartments (Figure 4D), Lamp1-positive vesicles (Figure 4E) and enzymatically active lysosomes (cathepsin L-positive vesicles; Figure 4F) in close proximity to internalized bacteria compared to PBS controls. While OM-89 expanded Lamp1- and LysoTracker-positive compartments (Figure 3G, 4B), colocalization of intracellular UPEC with these markers was largely proportional to vesicle abundance and consistent with stochastic overlap (enrichment scores ≈1 or <1) (Figure 4D, 4E and SI Figure 4B, 4C). However, OM-89 significantly increased the enrichment of UPEC in proximity to cathepsin L-positive vesicles (enrichment score 1.25 in OM-89 vs 0.91 in PBS) (Figure 4F, SI Figure 4D), indicating enhanced routing of UPEC into proteolytically competent autolysosomes. Overall, OM-89-treated bladder epithelial cells exhibited significantly elevated cathepsin L levels in the presence of infection (SI Figure 4E), indicating that the highly abundant acidic and lysosomal compartments were functionally active. The observed lysosomal expansion, acidification (Figure 4B, 4C), and increased cathepsin L activity (Figure 4F) independently confirm the transcriptomic enrichment of lysosome-associated pathways.

Since lysosomal proteases such as cathepsins require an acidic environment for activation of their antimicrobial effects against intracellular bacteria (Szulc-Dąbrowska et al, 2020), we next aimed to abolish acidification with pharmacological inhibition. BafA1 (Mauvezin & Neufeld, 2015) collapsed the OM-89-driven enrichment of intracellular UPEC in proximity to cathepsin L-positive vesicles to PBS-control levels, demonstrating a clear acidification dependence (Figure 4G, SI Figure 4F). Overall, baf1 or chloroquine (CQ) (Mauthe et al, 2018) treatment reduced cathepsin activity to comparable levels in OM-89- and PBS-treated cells (SI Figure 4E). Interestingly, CQ treatment led to a partial increase in cathepsin activity, particularly in infected PBS-treated cells.

Finally, given the enhanced lysosomal function and increased localization of intracellular UPEC to protease active compartments during OM-89 treatment, we next asked whether this effect was essential for OM-89-mediated protection. Importantly, inhibition of acidification using either bafilomycin A1 or chloroquine did not affect initial bacterial load during infection in OM-89-treated versus PBS-treated organoids (SI Figure 4G, 4H). However, blocking acidification using either of the two inhibitors fully abolished OM-89’s protective effect, restoring bacterial regrowth to levels observed in PBS-treated controls (Figure 4H, 4I; 10h pi – regrowth). Interestingly, while OM-89 still promoted antibiotic-mediated killing in bafilomycin A1-treated organoids (Figure 4H, killing), this effect was completely lost with chloroquine treatment (Figure 4I, 7h pi – killing).

Together, these data establish lysosomal acidification and protease activity as essential for the OM-89-dependent enhancement of intracellular bacterial clearance.

### OM-89 enhances intracellular activity across antimicrobial classes and UPEC strains

To validate OM-89’s ability to reduce regrowth of CFT073 after clinically relevant antibiotic exposure, we investigated its effect following treatment with antibiotics that have different mechanisms of action than ampicillin and different intracellular accumulation levels (Carryn et al, 2003; Bongers et al, 2019). We selected two commonly prescribed antibiotics for uncomplicated UTIs: Fosfomycin (FOS), a bacterial cell wall-targeting antibiotic displaying weak host cell accumulation (Höger et al, 1985) and trimethoprim/sulfamethoxazole (TMP-SMX), a folate synthesis inhibitor showing more effective intracellular accumulation (Bongers et al, 2019). OM-89 effectively reduced regrowth of CFT073 in bladder organoids (10h pi) after both FOS (Figure 5A) and TMP-SMX (Figure 5B) treatment and enhanced the killing (7h pi), indicating that its effect on bladder epithelial cells is independent of the antibiotic’s mode of action.

**Figure 5.**
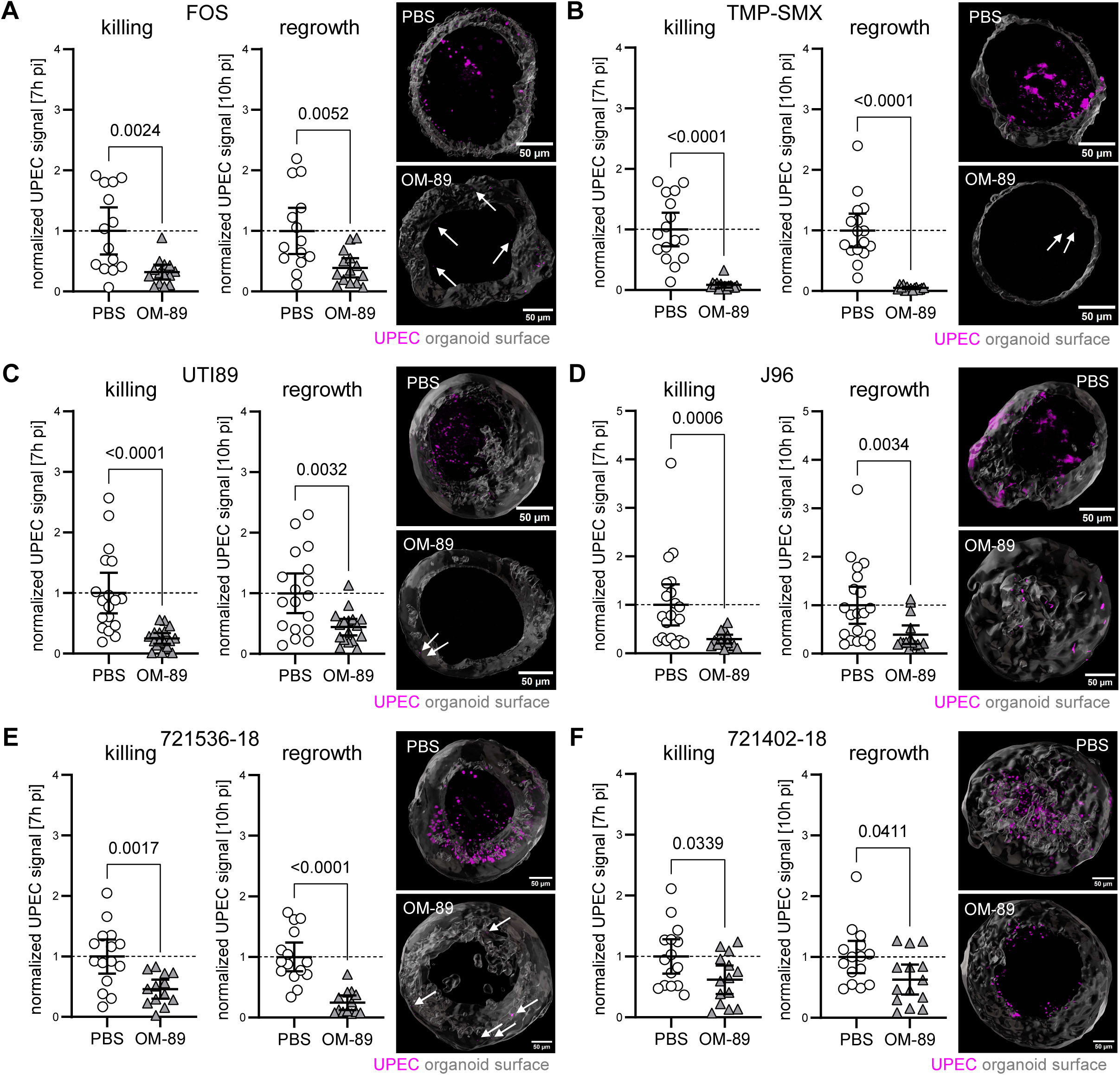
OM-89 reduces UPEC recurrence in bladder organoids across antibiotics and strains. (A), (B) CFT073 signal during treatment with (A) Fosfomycin (FOS) or (B) trimethoprim-sulfamethoxazole (TMP-SMX). Quantification of bacterial fluorescence area in mouse bladder organoids 7h pi (killing) and 10h pi (regrowth). Each dot represents one organoid. Values normalized to PBS control (dashed line). Mean ± 95% CI. Mann-Whitney test for killing in (B); Welch’s t test for (A) and regrowth in (B). N ≥ 14 per condition for (A) and n ≥ 15 per condition for (B). Representative organoids shown (grey, surface reconstruction) with UPEC (magenta, some marked with arrows). (C)-(F) Regrowth of (C) UTI89, (D) J96, (E) 721536-18 and (F) 721402-18 after ampicillin treatment (10x strain-specific MIC). Quantification of bacterial fluorescence area 7h pi (killing) and 10h pi (regrowth). Each dot represents one organoid. Values normalized to PBS control (dashed line). Mean ± 95% CI. Welch’s t test for killing in (E), (F) and regrowth in (C); Mann-Whitney test for killing in (C), (D) and regrowth in (D), (E), (F). N ≥ 17 per condition for (C), n ≥ 15 per condition for (D), n ≥ 13 per condition for (E) and n ≥ 14 per condition for (F). Representative organoids shown (grey, surface reconstruction) with UPEC (magenta, some marked with arrows).

Next, we assessed the regrowth of two additional well-characterized UPEC strains of phylogroup B2, UTI89 (a cystitis isolate) (Chen et al, 2006) and J96 (a pyelonephritis isolate) (Klein & Gitai, 2013), following ampicillin treatment. As observed with CFT073, OM-89 significantly enhanced killing (7h pi) of both UTI89 (Figure 5C) and J96 (Figure 5D) leading to reduced regrowth (10h pi), without affecting their initial growth within the organoids (SI Figure 5A, 5B).

Finally, we extended our analysis to two ampicillin-sensitive primary UPEC isolates obtained from cystitis patients, 721536-18 and 721402-18, which were identified among hyper-invasive strains in a genome-wide association study and belong to phylogroups A and B1, respectively (Cuénod et al, 2023). Similar to the laboratory strains, OM-89 did not affect initial bacterial loads within organoids (SI Figure 5C, 5D), indicating no direct antibacterial activity. However, during antibiotic treatment, OM-89 significantly enhanced killing (7h pi; Figure 5E, 5F) and reduced post-treatment regrowth of both primary isolates 721536-18 and 721402-18 (10h pi; Figure 5E, 5F), demonstrating that the positive effects of OM-89 extend beyond the classical B2 UPEC phylogroup to genetically and phylogenetically distinct clinical strains.

### OM-89 enhances lysosome-associated vesicle remodeling, acidification and protease activity in human bladder epithelial cells

The bladder epithelium differs in substantial ways between mouse and human – not only in the number of cell layers and the expression of specific markers, such as cytokeratin (CK)5, CK13, CK14, CK17 and toll like receptor (TLR)11 (Murray et al, 2021), but also cell-intrinsically for example in pathogen defenses (Holly et al, 2020; Zschaler et al, 2014). Such species-species factors could influence the outcome of infection and treatment processes.

Therefore, we asked whether the key hallmarks observed by the OM-89-driven phenotype, such as lysosome-associated vesicle remodeling, acidification, protease activation and antibiotic uptake observed in mouse bladder epithelium, are conserved in human cells. Using differentiated monolayers of primary human bladder epithelial cells (SI Figure 6), we quantified canonical autophagy and endo-lysosomal markers during infection and OM-89 treatment. OM-89 did not significantly change the number of p62- or LC3b-positive vesicles but significantly increased their size (SI Figure 7A, 7B), indicating formation of enlarged autophagosomal structures. In contrast to mouse bladder epithelial cells, OM-89 did not restore autophagic flux in human cells (SI Figure 7C), despite the significant increase in LC3b vesicle size.

**Figure 6.**
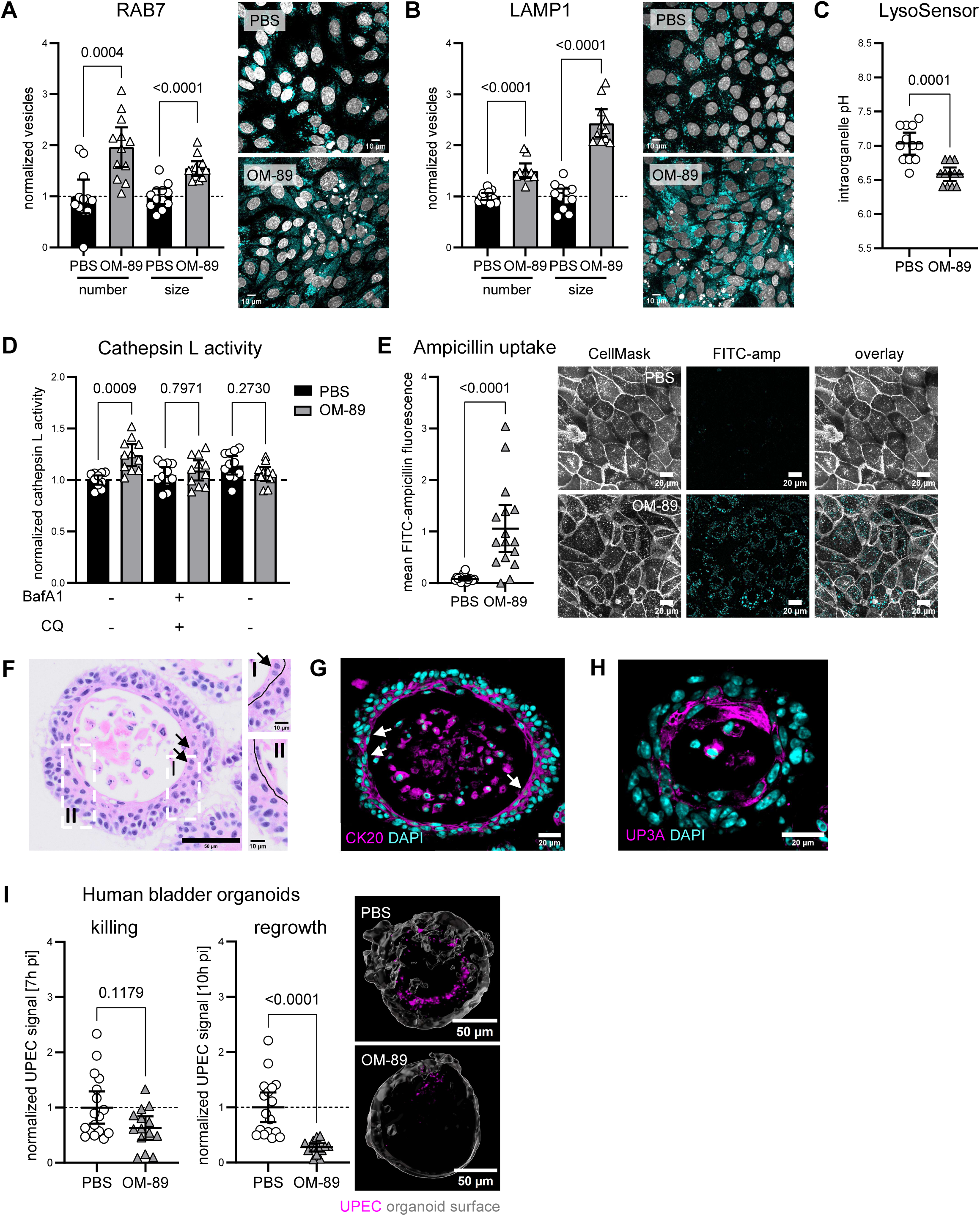
OM-89 enhances lysosomal activity, intracellular acidification, antibiotic uptake and bacterial clearance in human bladder epithelial systems. (A), (B) Quantification of vesicle numbers and size for (A) RAB7 and (B) LAMP1 in infected monolayers of human bladder epithelial cells with and without OM-89 treatment. Values normalized to PBS control (dashed line). Mean ± 95% CI. Welch’s t test for RAB7 (A); Mann-Whitney test for LAMP1 (B). N = 12 per condition for RAB7 and ≥ 11 per condition for LAMP1. Z-projection (maximum intensity) of representative images. Foci, cyan; DAPI, grey. (C) Intraorganelle pH in infected human monolayers with and without OM-89 treatment. Mean ± 95% CI. Mann-Whitney test. N = 12 per condition. (D) Cathepsin L activity in human monolayers during infection with or without OM-89, and with BafA1 or CQ blocking. Values normalized to unblocked PBS controls (dashed line). Mean ± 95% CI. Brown-Forsythe ANOVA with Welch’s correction. N = 11 per condition. (E) Uptake of FITC-labelled ampicillin into monolayers of human bladder epithelial cells. Quantification of fluorescence after background subtraction of PBS- or OM-89-treated cells without labelled antibiotic. Each dot represents one field of view. Mean ± 95% CI. Mann- Whitney test. N = 16 per condition. Z-projection (maximum intensity) of representative images. FITC-ampicillin shown in cyan. (F) Representative H&E (Hematoxylin & Eosin) staining of human bladder organoids derived from primary H-6215 cells, showing stratified epithelial architecture with 4-5 layers and thin, elongated cells lining the central lumen. Frequently binucleated cells are indicated by arrows. Scale bare, 50 μm. Inserts (I and II) highlight luminal cell morphology; black lines outline elongated umbrella-like cells. Scale bare, 10 μm. (G), (H) Human bladder organoids express markers specific to umbrella cells, cytokeratin (CK)20 (G), and important for UPEC infection, UP3A (H), towards the lumen. Binucleated cells are marked by asterisks. Representative images shown (partially same images as in SI Figure 9A). (I) CFT073 signal 7h pi (killing) and 10 h pi (regrowth) in human bladder organoids. Each dot represents one organoid. Values normalized to PBS control (dashed line). Mean ± 95% CI. Mann-Whitney test for killing and Welch’s t-test for regrowth. N ≥ 14 per condition. Images show representative organoids (grey, surface reconstruction) and UPEC (magenta, some marked with arrows).

We next assessed late endosomal and lysosomal compartments. OM-89 treatment significantly increased both the number and size of Rab7-positive vesicles and Lamp1-positive vesicles (Figure 6A, 6B), consistent with expansion and remodeling of the late endosomal-lysosomal network. These structural changes were accompanied by functional alterations: OM-89 significantly decreased intraorganellar pH (Figure 6C) and increased cathepsin L activity (Figure 6D), consistent with lysosome-centered functional activation. The increase in protease activity was fully reversible upon inhibition of acidification with bafilomycin A1 or chloroquine (Figure 6D), demonstrating dependence on lysosomal acidification. Finally, OM-89 markedly enhanced intracellular uptake of fluorescently labeled ampicillin in human bladder epithelial cells (Figure 6E), mirroring the uptake phenotype observed in mouse cells.

Together, these data show that OM-89 drives conserved lysosome-centered remodeling and enhanced antibiotic uptake in human bladder epithelium, while revealing species-specific differences in autophagic flux regulation.

### OM-89 enhances antibiotic-mediated killing and limits UPEC regrowth in human bladder organoids

Finally, we asked whether OM-89 exerts similar effects on bacterial clearance in human bladder epithelium. We therefore established human bladder organoids (hBOs) from primary bladder epithelial cells which already showed robust stratification and differentiation in another 3D microtissue model (Paduthol et al, 2026). Cells were cultured in 3D under conditions adapted from the mouse bladder organoid model (Sharma et al, 2021) and other bladder epithelial models (Russell et al, 2023). Briefly, following an initial expansion phase of 9 days, differentiation was induced for 4-5 days to generate stratified organoids that recapitulated key aspects of human urothelium (SI Figure 8A, 8B). By day 13-14, hBOs displayed 3-4 epithelial layers surrounding a central lumen (Figure 6F, SI Figure 9A). Luminal cells exhibited hallmark features of umbrella cells, including thin morphology, frequent binucleation, and expression of CK20+ and UP3A+ cells (Khandelwal et al, 2009; Wu et al, 2009) (Figure 6G, 6H, SI Figure 9A), while CK13+ intermediate and CK5+ basal-like cells (Jafari & Rohn, 2022) populated deeper layers (SI Figure 9A). RT-qPCR confirmed upregulation of umbrella (*KRT20*, *UPK3A*) and intermediate/progenitor (*KRT13*, *TP63* (Pignon et al, 2013)) markers (SI Figure 9B). The organoids formed a functional epithelial barrier, as evidenced by *ZO-1* expression (Smith et al, 2015) and restricted dextran diffusion in both freshly cultured and cryopreserved-then-differentiated (thawed) organoids (SI Figure 9C, 9D).

To test OM-89’s effect on human cells, hBOs were treated under the continuous application regimen for 72 hours prior to microinjection of UPEC strain CFT073, with OM-89 maintained throughout the experiment. As in mouse organoids, OM-89 did not alter bacterial burden during the initial infection phase (4h pi; SI Figure 10A). However, although OM-89 did not significantly enhanced antibiotic-mediated killing in hBOs (7h pi; 10x MIC ampicillin), it significantly reduced bacterial regrowth following antibiotic treatment (10h pi) (Figure 6I). Consistent with our observations in mBOs, UPEC regrowth in hBOs preferentially originated from the organoid wall rather than from luminal compartments (SI Figure 10B, SI movie 3 and 4), indicating that the epithelial layer serves as the primary site of bacterial regrowth following antibiotic withdrawal.

Together, these findings demonstrate that OM-89 restricts intracellular UPEC regrowth in a human organoid model, suggesting conservation of its positive effects across host species.

## Discussion

Intracellular reservoirs of UPEC within bladder epithelial cells are a major driver for the reactivation of UTIs from the bladder epithelium due to their capacity to evade both antibiotics and immune responses (Mysorekar & Hultgren, 2006). Here, we identify a host-directed approach that strengthens epithelial antimicrobial mechanisms and improves intracellular antibiotic efficacy. Using the bacterial lysate OM-89 – a clinically approved therapy for recurrent UTIs with decades of clinical use – as a mechanistic probe of epithelial responses, we show that bladder epithelial cells can be modulated to accelerate intracellular bacterial clearance through coordinated remodeling of lysosomal pathways and enhanced intracellular accessibility of antibiotics. Transcriptomic profiling further revealed that OM-89 induces a transcriptional program partially overlapping with infection-associated responses, yet remaining clearly distinct from the infected state, suggesting that OM-89 establishes a modified epithelial defense state rather than mimicking infection-driven inflammation. Together, these findings provide a mechanistic framework for the long-observed clinical efficacy of OM-89. Our findings reveal that the urothelium itself can be therapeutically targeted to reduce pathogen regrowth by transforming the epithelial barrier from a passive refuge for UPEC into an active defense site.

OM-89 induced coordinated lysosome-centered functional remodeling in both mouse and human bladder epithelial cells. In mouse cells, the observed increased Rab11A and Rab27B vesicles are consistent with enhanced endosomal turnover and secretion pathways implicated in expulsion of intracellular pathogens (Miao et al, 2017). Despite accumulation of p62, likely reflecting infection-associated cellular stress (Kumar et al, 2022), enlarged autophagosomes together with increased autophagic flux indicate that the ubiquitinated cargo is efficiently routed toward lysosomal degradation in mouse epithelial cells, overcoming typical UPEC-mediated blockade of autophagic maturation (Li et al, 2024). Increased numbers of Rab7-positive late endosomes and Lamp1-positive vesicles further support expanded lysosomal biogenesis and remodeling. While OM-89-induced lysosomal remodeling, autophagic vesicle enlargement and acidification were also preserved in human bladder epithelial cells, the effect on canonical autophagic flux appears to diverge, with restoration of flux in mouse but not in human cells. One underlying reason could be the difference in basal levels of autophagy between the two species, as at least mouse models seem to have generally a higher basal level of autophagy than human lung epithelial cells (Zhao et al, 2019). Such species-specific variability likely reflects differences in regulatory wiring upstream of autophagy, leading to differences in the tuneability of these networks by OM-89, but suggest that the core lysosome-directed antimicrobial mechanism appears to be preserved even when certain pathway features differ between species.

Strikingly, OM-89 did not only enlarge the lysosomal network but enhanced its functional capacity in both species. Lysosomal proteases became concentrated in proximity to intracellular pathogens, suggesting targeted trafficking of UPEC into proteolytically active autolysosomes. Pharmacological inhibition of lysosomal acidification abolished OM-89-mediated protection, demonstrating a direct link between lysosomal function and reduced UPEC regrowth. These coordinated changes likely undermine UPEC’s strategies to evade degradation (Naskar et al, 2023; Miao et al, 2015), revealing a yet underappreciated mechanism by which OM-89 fine-tunes epithelial antimicrobial effects via lysosomal activation.

Although this enhanced host activity did not affect UPEC replication during primary infection, its significance emerged during antibiotic treatment as observed already in a mouse study using repeated UPEC infections (Canton et al, 2025). Using bladder organoid and monolayers models, we showed that OM-89 increased intracellular uptake and/or retention of structurally distinct, poorly permeant antibiotics, as well as dextran, indicating a broad effect on intracellular delivery pathways. Because antibiotics used to treat UTIs display varied intracellular activity profiles (Carryn et al, 2003; Höger et al, 1985; Bongers et al, 2019), improving intracellular access may significantly expand their therapeutic efficacy. Supporting this, OM-89 reduced bacterial regrowth even at low antibiotic concentrations, indicating that host-directed boosting of intracellular pharmacokinetics can enhance antibiotic performance.

OM-89’s protective effect extended across multiple antibiotics and genetically diverse UPEC strains – from classical UTI-associated B2 phylogroup isolates to phylogroups A and B1, more commonly linked to gut strains (Wang et al, 2023) – indicating robustness against heterogeneous intracellular persistence strategies. Importantly, these epithelial effects were not restricted to murine systems. Human bladder organoids recapitulated the functional phenotype, showing enhanced antibiotic-mediated killing and reduced post-treatment regrowth. Together with the conserved core lysosome-centered antimicrobial response, our findings support the translational relevance of OM-89 as a clinically used therapy whose mechanism of action involves direct modulation of epithelial antimicrobial pathways.

Although a clear effect of OM-89 on spontaneous recurrences and bacteriuria was demonstrated in mice previously, the study did not find depletion of UPEC from the bladder reservoir (Canton et al, 2025). Our findings suggest that enhanced endo-lysosomal trafficking of intracellular UPEC together with increased acidification may restrict bacterial reactivation and therefore luminal shedding. Such modulation of epithelial trafficking of intracellular pathogens could therefore mechanistically contribute to reduced recurrences of UTIs by limiting the epithelial source of reseeding events. Such host-targeted strategies at the site of infection, the bladder epithelium, could enhance the effectiveness of existing antibiotics at the cellular level and help limit intracellular pathogen survival, shorten treatment courses, and reduce overall antibiotic exposure. In the face of rising antimicrobial resistance (2024), strengthening epithelial antimicrobial function offers a complementary route to shift the bladder mucosa from a passive niche of bacterial survival and infection reactivation toward an active site of accelerated pathogen clearance.

## Methods

### Bacteria

The well-characterized UPEC strains CFT073 (Mobley et al, 1990), UTI89 (Chen et al, 2006) and J96 (Klein & Gitai, 2013), and primary clinical isolates 721536-18 and 721402-18 (Cuénod et al, 2023) were used. Strains were grown overnight in DMEM/F-12 (Thermo Fisher) supplemented with 20 mM HEPES (Thermo Fisher) without (for organoid injections) or with (for monolayer infections) 10% heat-inactivated Fetal Bovine Serum (FBS, Thermo Fisher) at 37°C statically.

#### Chromosomal sfGFP integration

To enable real-time monitoring of the infection dynamics, all UPEC strains were genetically engineered to stably express *sfGFP* under control of a strong P_σ70_ promoter at the chromosomal *att_HK022_* site using scare-less cloning (Eshaghi et al, 2016; Simonet et al, 2022). Briefly, the *kan^R^*-P_RhaB_-*relE* toxin cassette was amplified from plasmid pSLC217 using primers P4 and P5 and integrated into the genome of the strains using λ Red recombineering (Datsenko & Wanner, 2000). Primers P6 and P7 were then used to amplify the sfGFP-containing fragment from plasmid pSLC293, and these PCR amplicons were used to replace the *kan^R^*-P_RhaB_-*relE* toxin cassette by λ Red recombineering with negative selection on M9 agar (Sigma) supplemented with 2% L-rhamnose (Sigma). Successful recombination at each step was confirmed by PCR using primers attHK_F and attHK_R.

Used bacterial strains are listed in Table 1. Primers are listed in Table 2.

**Table 1.** Used strains.

| Strain name | Genotype | Source |
| --- | --- | --- |
| TS4 | CFT073 $\Delta fliC::FRT$ | Simonet et al. 2022 |
| TS85 | CFT073 *att<sub>HK022</sub>::$\sigma_{70}$-sfGFP* | Simonet et al. 2022 |
| TS66 | TS4 *att<sub>HK022</sub>::$\sigma_{70}$-sfGFP* | Simonet et al. 2022 |
| FN27 | J96 *att<sub>HK022</sub>::$\sigma_{70}$-sfGFP* | Valentin Borgeat |
| FN28 | UTI89 *att<sub>HK022</sub>::$\sigma_{70}$-sfGFP* | Valentin Borgeat |
| VB36 | 721536-18 *att<sub>HK022</sub>::$\sigma_{70}$-sfGFP* | this study |
| VB37 | 721402-18 *att<sub>HK022</sub>::$\sigma_{70}$-sfGFP* | this study |

**Table 2.**
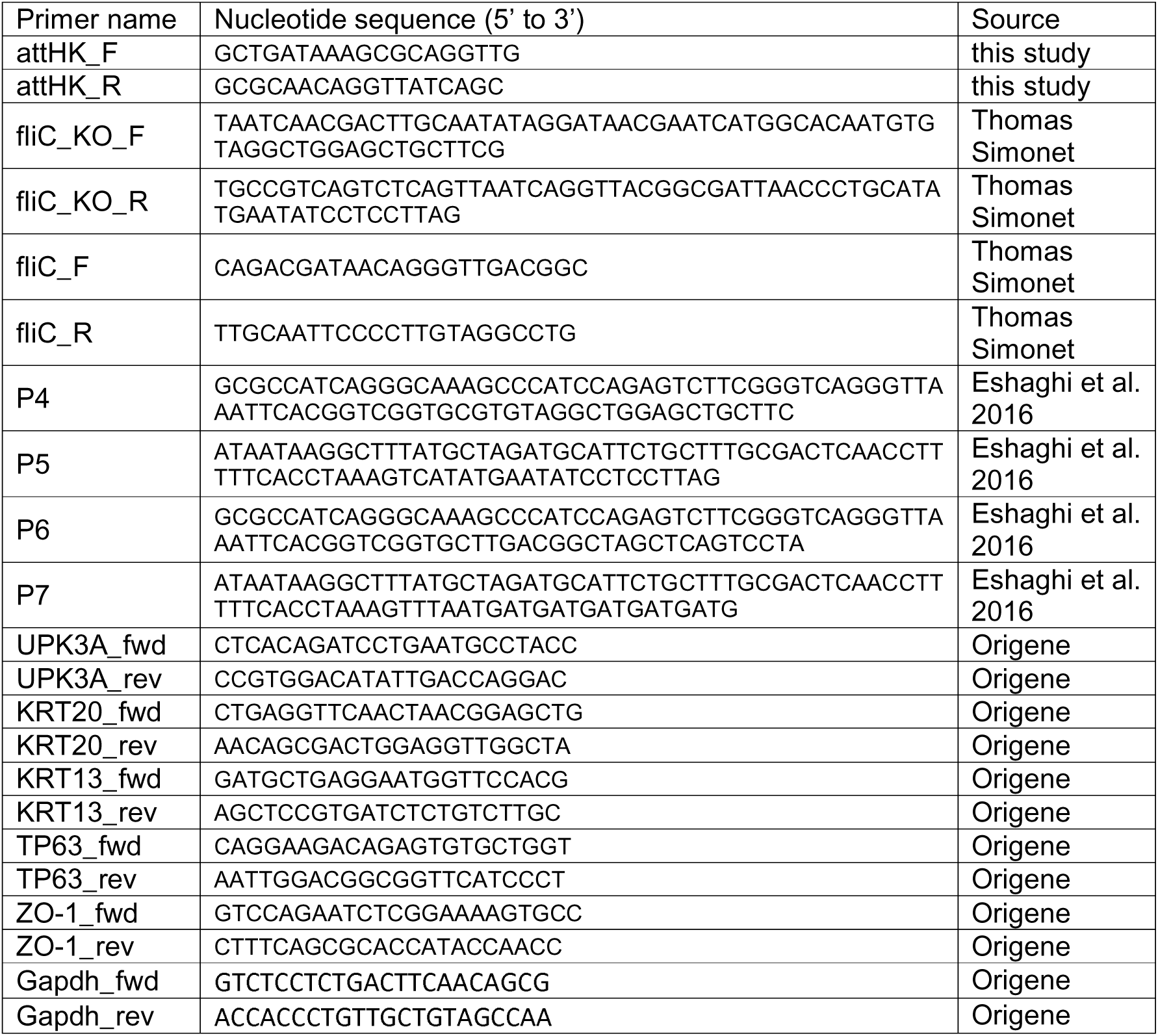
Used primers.

### Cell culture

Mouse primary bladder epithelial cells were obtained from C57BL/6 mice (Charles River Laboratories) aged 3-4 months or female ROSA^mT/mG^ mice (Jackson Laboratories) at age 4 months. Mice were housed in a specific pathogen-free facility. All animal protocols were reviewed and approved by EPFL’s Chief Veterinarian, by the Service de la Consommation et des Affaires Vétérinaires of the Canton of Vaud, and by the Swiss Office Vétérinaire Fédéral.

Human primary bladder epithelial cells (H-6215, CellBiologics) were cultured in H-6621 medium (CellBiologics) supplemented to a final concentration of 10% heat-inactivated fetal bovine serum (FBS) (Thermo Fisher). Cells were received from the supplier at passage 3 and expanded according to the manufacturer’s recommendations before use at passages 6-8 (maximum passage as recommended by the supplier).

Routinely, mouse and human cells were grown at 37°C and 5% CO_2_ without antimicrobial or antimycotic agents. Organoids were cultured in media without FBS but with 2.5 mM GlutaMAX (Thermo Fisher) supplementation, whereas for monolayers a total of 10% heat-inactivated FBS was added and plastics were coated with gelatin-based coating solution (ABC Biopply). The day prior to experiments, either medium was replaced with DMEM/F12 (without phenol red) supplemented with 20 mM of HEPES (Thermo Fisher).

#### Mouse bladder organoids

Mouse bladder organoids (mBOs) were generated as previously described (Sharma et al, 2021) with minor modifications.

Briefly, excised mouse bladders were minced with scissors and incubated in 1-1.5 ml pre-warmed TryPLE (Thermo Fisher) at 37°C with shaking at 170 rpm for 30 min. Following vortexing for 30 s, TryPLE was neutralized by addition of DMEM/F-12 supplemented with 10% FBS. Single cells were embedded in reduced growth factor basement membrane extract (BME; BioTech AG) and seeded as 50 µl domes in 24-well plates in an inverted configuration for 30 min at 37°C to promote 3D growth. Thereafter, organoids were fed with mouse bladder organoid (MBO) medium (Advanced DMEM/F-12 medium (Thermo Fisher) supplemented with 2.5 mM GlutaMAX (Thermo Fisher), 100 ng/ml FGF10 (Peprotech), 100 ng/ml FGF7 (Peprotech), 500 nM A83-01 (Tocris Bioscience), 2% B27 (Thermo Fisher), and 10 µM Y-27632 ROCK inhibitor (Abmole Bioscience) and 100 μg/ml primocin (InvivoGen). Typically, roughly 50,000 bladder cells were seeded per dome as not all of them have the necessary stem-cell like characters to promote growth of organoids. MBO medium was renewed every 2-3 days and upon first splitting or passaging, primocin was omitted.

When organoids became overly dense, they were split into multiple BME domes following extraction using 250 µl ice-cold Cell Recovery Solution (Corning) per dome and incubating for 30 min on ice. After centrifugation at 300 g for 5 min, the supernatant was carefully removed and the organoids in remaining BME were washed once in ice-cold PBS before spinning again. Grown organoids were passaged every 5-7 days by extraction from BME with Cell Recovery Solution, followed by TryPLE digestion for 10 min at 37°C to generate single cells. At this stage, roughly 5,000-10,000 cells were seeded per dome. For cryopreservation, culture medium was removed and organoids embedded in BME were directly submerged in freezing medium (60% FBS, 30% HBO medium, 10% DMSO) prior to freezing.

tdTomato-expressing organoids derived from ROSA^mT/mG^ mice were used for microinjection experiments, whereas wild-type (WT) organoids dissociated into single cells were primarily used to grow as monolayers.

Generally, all plastics, except well plates, were pre-coated with 1% bovine serum albumin (BSA) in PBS to avoid organoid loss due to increased adhesion of organoids to uncoated plastic.

#### Human bladder organoids

Human bladder organoids (hBOs) were generated from H-6215 cells expanded as described above. Single cells at passages 6-8 were seeded at a density of ≤ 5,000 cells per 50 µl BME dome in 24-well plates. As for mouse bladder organoids, laboratory plastic was pre-coated with 1% BSA in PBS and cells were seeded in an inverted configuration for 30 min at 37°C to promote 3D growth.

Organoids were cultured for 9 days in a 1:1 mixture of HBO medium (MBO medium as described above, supplemented with 1 mM N-acetyl L-cysteine (NAC) (Sigma), 10 mM Nicotinamide (NIC) (Sigma) and 50 ng/ml EGF (Thermo Fisher)) and LWRN-conditioned medium (Sigma) (Russell et al, 2023), referred to as 50/50 medium. Medium was refreshed every 2 days. On day 9, organoids were either differentiated or cryopreserved.

For cryopreservation, culture medium was removed and organoids embedded in BME were directly submerged in freezing medium (60% FBS, 30% HBO medium, 10% DMSO) prior to freezing. Following thawing, organoids were re-embedded in 50 µl BME domes and allowed to recover overnight in 50/50 medium before differentiation.

Differentiation was induced by culturing organoids for 4-5 days in 95% HBO medium supplemented with 5% LWRN-conditioned medium (95/5 medium), with medium changes every 2 days. Differentiated hBOs were used for characterization or microinjection experiments on days 13-14.

#### Mouse and human bladder epithelial cell monolayers

Single cells of mouse bladder epithelial cells were obtained by dissociating WT mBOs using TryPLE digestion for 10 min at 37°C. Single-cell suspensions of mouse bladder epithelial cells or human bladder epithelial cells (H-6215, passages 6-8) were seeded into gelatin-coated µ-Plate 96-well dishes (ibidi) at densities of 1.5-2*10L cells per well and 1-1.5*10L cells per well, respectively. Cells were cultured in 200 µl MBO medium or H-6621 medium supplemented with 10% FBS, respectively. Upon reaching approximately 40-50% confluency (typically the following day), the respective culture media were replaced with DMEM/F-12 supplemented with 10% FBS and cells were maintained under these conditions for 3 days to promote epithelial differentiation through growth factor withdrawal. Differentiated monolayers were used for experiments upon reaching 100% confluency. Umbrella-like cells expressed cytokeratin (CK)20 and uroplakin (UP)3a, intermediate cells expressed CK13, basal cells expressed CK5 (Jafari & Rohn, 2022; Sharma et al, 2021). Umbrella-like cells were frequently binucleated.

### OM-89 treatment

Organoids and monolayers were treated with OM-89 (OM-Pharma) at 500 µg/ml of protein in either MBO, 95/5 HBO/LWRN media or DMEM/F-12 media supplemented with 2.5 mM GlutaMAX and 20 mM HEPES. For control groups, equal volumes of phosphate buffered saline (PBS, Thermo Fisher) was used instead of OM-89. OM-89 and PBS were added to the media according to the different treatment regimens: continuous application – 72 hour treatment pre-infection and treatment throughout the experiment, pre-application – 72 hour treatment pre-infection only with a 24 hour resting period where treatment was omitted, co-application – treatment only during antibiotic treatment. For the continuous application, bacteria were microinjected after adjusting their optical density at 600 nm (OD_600_) in media with either PBS or OM-89. If not stated other, bladder epithelial cells were subjected to the continuous application.

### Minimal inhibitory concentration (MIC) measurements

Bacteria were grown in DMEM/F-12 media (without phenol red) supplemented with with 2.5 mM GlutaMAX and 20 mM HEPES. Overnight cultures were diluted to an OD_600_ of 0.002 and grown till OD_600_ of 0.01. Bacterial cultures were adjusted to an OD_600_ of 0.0005 for serial dilutions of tested antibiotics. MIC was estimated based on growth after 24 hours in a plate reader (Tecan Infinite M Nano) and using the Goempertz equation for MIC determination (Lambert & Pearson, 2000) (Table 3). If not stated other, 10x the strain MIC of the indicated antibiotic was used for infection experiments.

**Table 3.** MICs.

| Strain | Antibiotic | MIC [µg/ml] |
| --- | --- | --- |
| CFT073 | Ampicillin | 6.43 |
| CFT073 | Fosfomycin | 6.97 |
| CFT073 | Trimethoprim/Sulfamethoxazole | 1.76 |
| UTI89 | Ampicillin | 5.35 |
| J96 | Ampicillin | 9.98 |
| 721536-18 | Ampicillin | 4.98 |
| 721402-18 | Ampicillin | 11.61 |

### Injection experiments

Organoid injections were performed as described (Sharma et al, 2021) with minor modifications.

Briefly, the day prior to the experiments, organoids were extracted from BME domes using cell recovery solution and transferred to µ-Slide 2 Well dishes (ibidi) to allow direct comparison of OM-89-treated and PBS-treated conditions. Organoids were embedded in 75 µl BME spread over an area of approximately 1.5 cm^2^ to align organoids in a narrow focal plane rather than a dome configuration. At this stage, the media was replaced by phenol red-free DMEM/F-12 medium with 2.5 mM GlutaMAX and 20 mM HEPES with either OM-89 or PBS.

On the day of the microinjections, hBOs were stained inside the BME with CellTracker™ Red CMTPX (Thermo Fisher) at 5 µM for 1 hours, whereas mBOs derived from ROSA^mT/mG^ mice were left unstained due to their inherent tdTomato-expression. CellTracker was washed off with excessive volumes of warm DMEM/F-12, before putting again phenol red-free DMEM/F-12 medium with 2.5 mM GlutaMAX and 20 mM HEPES with either OM-89 or PBS.

#### Bacterial injection

UPEC strains with stable chromosomal integration of *sfGFP* were grown overnight statically at 37°C in DMEM/F-12 (phenol red-free) media with 2.5 mM GlutaMAX and 20 mM HEPES.

To enable media exchange during live imaging, a custom-made perfusion lid compatible with µ-Slide 2 Well dishes was fabricated the day before microinjections with one inlet and one outlet per well. Using a drill press, two small holes were drilled at the outer corners of a DIC lids for µ-Slides (ibidi) in each well to fit cannula tips with conical, tapered ends (1.30/0.75 x 15 mm, Unimed). Cannula tips were connected to roughly 20 cm of silicone tubing (ELASTOSIL® R plus 4305/60, Freudenberg). Each cannula tip was secured in place with glue with minimal distance to the bottom of the dish to allow easy media change without leaving a dead volume. The tubing was split mid-length with additional cannula connectors to allow easy routing into an H201-K-Frame incubation chamber (Okolab). At the distal end, the silicone tubing was connected to dispensing tips (LL 1/2”, ID 0.69 mm, Gonado) attached to 10 ml syringes for manual media exchange. For every media exchange (e.g. antibiotic treatment or withdrawal), each well was washed with 9 ml of the respective media before leaving 1 ml in the well. All lid components were disassembled and cleaned with ethanol between experiments, whereas silicone tubing was replaced after each use.

In the morning of the experiments, the media of the organoids was replaced with fresh phenol red-free DMEM/F-12 supplemented with 2.5 mM GlutaMAX and 20 mM HEPES with either OM-89 or PBS. The OD_600_ of bacterial cultures was adjusted to 2.25 in a 1:1 dilution of media and Phenol Red solution (Sigma-Aldrich) to facilitate the visualization of injected organoids. Microinjections were performed using a Pneumatic PicoPump (WPI) using micropipettes prepared from thin wall glass capillary (TW100F-4 with length 100 mm and diameter 1 mm) using a Flaming/Brown Micropipette Puller (Sutter Instruments model P-87) set at pressure 360, heat 866, Vel 200. The micropipettes were then cut with tweezers under a stereomicroscope (Olympus SZX-16) to generate a tip size ejecting roughly 1 nl volume of the adjusted bacterial culture in mineral oil on a Zeiss coverslide (corresponding to 100 mm), corresponding to 1926 ± 150 colony forming units (CFU) per injection (quantified from two needles, plating three replicates on LB agar plates). Injected organoids were allowed to rest for 1 hour prior to time-lapse microscopy imaging onset after assembling the µ-Slide 2 Well dishes with the custom-made perfusion lid.

#### Dextran injection of hBOs

hBOs were microinjected with fluorescein isothiocyanate (FITC)-tagged dextran (4000 Da, Sigma Aldrich) at a final concentration of 1 mg/ml after diluting dextran dissolved in DMEM/F-12 (without phenol red) supplemented with 2.5 mM GlutaMAX and 20 mM HEPES 1:1 with Phenol Red solution. Dextran diffusion out of the organoids was monitored for 2 hours post-injection.

### Time-lapse microscopy

Imaging of infected organoids was conducted using a Leica Thunder DMi8 microscope (Leica) with a temperature-controlled microscope environmental chamber maintained at 37°C and 5% CO_2_ in a stage-top chamber (OKOlabs) using a Leica HC FLUOTAR VISIR 25x (NA 0.95) water-immersion objective. To maintain the water immersion for the objective, water was pumped to the ring around the water objective. Microinjected organoids were identified and imaged at ex476/em519 nm (UPEC) and ex554/em594 nm (organoids) at 85 ms exposure time, acquiring the multiple channels during the same imaging sequence to improve the temporal resolution. Z stacks were acquired with 1 µm step sizes. Selected organoids were images 4 hours into the infection phase (corresponding to G3 of acquired images), throughout the antibiotic phase where indicated (corresponding to Amp0-3 of acquired images), straight after antibiotic removal (corresponding to RG0 of acquired images) and 3 hours after antibiotic withdrawal (corresponding to RG3 of acquired images). For experiments indicated, antibiotic treatment or regrowth phases were extended. UPEC fluorescence area overlapping with organoid area (meaning bacterial signal only from inside the infected organoid by masking the organoid signal on the GFP signal; the intra-organoid bacterial signal) was extracted at the indicated time-points using fiji scripts *Volumetric calc_growth* and *Volumetric calc_regrowth*. For analysis, the obtained fluorescence area at any time-point (growth, antibiotic-mediated killing or regrowth) of the intra-organoid bacterial signal was normalized to the same time-point of the PBS control group within one experiment. The intra-organoid masking therefore includes all bacterial signal measured within the organoid; luminal bacteria and tissue-associated bacteria together.

### Blocking of acidification

Where indicated, 10 µM chloroquine (CQ, InvivoGen) or 0.1 µM bafilomycin A1 (bafA1, InvivoGen) was added for 3 hours prior to infection experiments. Afterwards, blocking was washed off with excessive amounts of prewarmed DMEM/F-12 before adding fresh media before UPEC infections. When enhancing for intracellular bacteria, such as co-localization experiments with Cathepsin L, blocking was added 1 hour before infection, and left throughout the infection and Cathepsin L staining (additional 4 hours).

### Immunofluorescence staining

#### Staining of paraffin-embedded human bladder organoids

hBOs at day 13-14 were extracted from BME with Cell Recovery Solution and fixed in 4% paraformaldehyde (PFA, Thermo Fisher) in PBS for 1 hour at 4°C. After washing, fixed organoids were resuspended in 50 µl of prewarmed Histogel (Thermo Fisher) at 50°C and pipetted out as a small hemispherical dome inside a 1-cm Tissue-Tek Cryomold. The cryomold was placed on a cold ice plate for solidification. Subsequently, the hemispherical Histogel was processed for paraffin embedding. Organoids embedded in paraffin were cut into 4 µm slices. The thin paraffin sections were deparaffinized and rehydrated by immersing the slides through the following solutions: xylene, three washes of 5 min each; 100% ethanol, two washes of 10 min each; 95% ethanol, two washes of 10 min each; 70% ethanol, two washes of 10 min each; 50% ethanol, two washes of 10 min each; PBS, three washes of 5 min each. Rehydrated slides were then processed for heat-induced antigen retrieval using 10 mM citrate buffer (pH 6.0). Slides were washed three times with PBS, permeabilized with 0.1% Triton X-100 for 15 min, washed three with PBS, and blocked with 1% BSA in PBS for 1 hour. The boundaries of paraffin sections were marked with a hydrophobic pen. Primary antibodies against cytokeratin (CK)7 for general bladder epithelium, CK5 for basal cells, CK13 for intermediate cells, and CK20 and uroplakin (UP)3a for umbrella cells were diluted 1:100 in 1% BSA in PBS and incubation was performed overnight at 4°C. After washing three times for 10 min each in PBS, slices were incubated with 1:1000 dilution of secondary antibody and DAPI in 1% BSA in PBS for 1 hour at room temperature. All used antibodies are listed in Table 4. Images were taken on a Confocal Stellaris 5 (Leica) using a HC PL APO 63x (NA 1.40) oil-immersion objective.

**Table 4.** Used antibodies.

| Antigen | Used reactivity | Host | Supplier | Catalog Number |
| --- | --- | --- | --- | --- |
| LC3b | mouse, human | rabbit | Abcam | ab192890 |
| Rab11A | mouse | mouse | Santa Cruz | sc-166912 |
| Rab27B | mouse | rabbit | Biorbyt | orb1264367 |
| Rab7 | mouse, human | mouse | Santa Cruz | sc-376362 |
| Lamp1 | mouse | rat | Santa Cruz | sc-19992 |
| Lamp1 | human | mouse | Abcam | ab25630 |
| p62 | mouse, human | rabbit | Abcam | ab109012 |
| UP3A | mouse | mouse | Santa Cruz | sc-166808 |
| UP3A | human | rabbit | Thermo Fisher | PA5-87581 |
| CK20 | mouse, human | rabbit | Biorbyt | orb256650 |
| CK13 | mouse, human | rabbit | Abcam | ab198584 |
| CK7 | human | rabbit | Abcam | ab181598 |
| CK5 | mouse, human | rabbit | Abcam | ab52635 |
| GAPDH | mouse, human | rabbit | Abcam | ab181602 |
| HRP | anti-rabbit IgG | goat | Agilent | P0448 |
| Alexa Fluor-488 | anti-rabbit IgG | donkey | Thermo Fisher | A-21206 |
| Alexa Fluor-647 | anti-rabbit IgG | donkey | Thermo Fisher | A-31573 |
| Alexa Fluor-647 | anti-mouse IgG | donkey | Thermo Fisher | A-31571 |

#### Differentiated monolayer stainings

Confluent mouse and human monolayers were fixed with 4% PFA for 1 hour at 4°C and then washed with PBS. For umbrella-like cells, cells were blocked for 1 hour at room temperature in 1% BSA before staining with UP3A in 1% BSA in PBS. After washing, cells were permeabilized with 0.1% Triton-X in PBS for 10 min. Cells were washed and blocked again before staining with primary antibodies for the main three bladder epithelial cell types: cytokeratin (CK)5 for basal cells, CK13 for intermediate cells and CK20 for umbrella-like cells. Primary antibodies were diluted 1:100 in 1% BSA in PBS. After washing, cells were incubated with 1:1000 dilution of secondary antibody, phalloidin-555 (Thermo Fisher) and DAPI in 1% BSA in PBS. Each staining was carried out for 1 hour at room temperature. All used antibodies are listed in Table 4. Images were taken on a Confocal Stellaris 5 using a HC PL APO 63x (NA 1.40) oil-immersion objective.

#### Vesicle staining

Differentiated monolayers were infected with TS4 at roughly 1*10^6^ CFU/ml (effective 1*10^5^ CFU/well/100 µl) for 4 hours. Thereafter, extracellular bacteria were washed with excessive amounts of PBS before fixing. Fixing, permeabilization and staining were carried out as mentioned above. All used antibodies are listed in Table 4. Images were taken on a Confocal Stellaris 5 using a HC PL APO 63x (NA 1.40) oil-immersion objective. Foci were counted in 3D using fiji script *3D_foci_counting*.

### Western blot

Proteins from differentiated and TS4 infected monolayers were isolated by resuspending cells after TryPLE digestion in 300 µl Radioimmunoprecipitation assay (RIPA) buffer. The protein mixture was then either used immediately or aliquoted and stored at -80°C. Protein concentrations were measured with the Pierce BCA protein assay kit (Thermo Fisher) and approximately 3 µg of proteins were boiled for 10 min at 70°C with NuPAGE LDS sample buffer (Invitrogen) and Dithiothreitol (DTT, Thermo Fisher) before running them on a 4-12% Bis-Tris gel (Invitrogen). Proteins were transferred onto a membrane using the iBlot Gel Transfer Device (Invotrgen). The membrane was cut to sperate GAPDH and LC3b bands and the membrane part containing LC3b (15-17 kDA) was pretreated with SuperSignal West Atto substrate solution (ThermFisher). After blocking both membrane parts for 1 hour at room temperature with 1% BSA in Tris-buffered saline (TBS), they were incubating with primary antibodies overnight at 4°C – GAPDH 1:2000 in 2% BSA in TRIS-NaCl-Tween 20 (TNT) buffer; LC3b 1:1000 in SuperSignal western blot enhancer (Thermo Fisher). After washing with TBS, blots were incubated with secondary antibody 1:2000 in 2% BSA in TNT for 1 hour at room temperature prior to visualizing signals with SuperSignal West Pico PLUS chemiluminescent substrate (Thermo Fisher). All used antibodies are listed in Table 4.

### Uptake assays

#### Dextran-TMR uptake

Differentiated mouse monolayers were incubated with fixable Dextran Tetramethylrhodamine (Dextran-TMR; Thermo Fisher) at a final concentration of 1 mg/ml for 4 hours before repeated washes with PBS and fixation with 4% PFA for 1 hour at 4°C. Images were taken on a Confocal Stellaris 5 using a HC PL APO 63x (NA 1.40) oil-immersion objective. To reduce low-frequency background variations, a Gaussian-blurred duplicate of the projection was subtracted from the original image. Background correction was performed using separate control images acquired under identical microscope settings in the absence of the labeled compound. Intracellular fluorescence was quantified after background subtraction of images of PBS or OM-89-treated cells using fiji script *mean_fluorescence*.

#### Labelled antibiotic uptake

Ampicillin and gentamycin were labelled using Pierce™ FITC Antibody Labeling Kit (Thermo Fisher) according to the manufacturer’s recommendations. Labelled stocks at 2 mg/ml of each antibiotic were aliquoted and stored at -20°C. FITC-labelling rendered the antibiotics ineffective at the used concentrations.

Differentiated monolayers were incubated with FITC-labelled ampicillin and gentamycin for 3 hours at a final concentration of 64.5 µg/ml (10x MIC of CFT073 for unlabelled ampicillin) and 100 µg/ml (usual concentration used for gentamicin protection assays (Mulvey et al, 2001)) respectively. Monolayers were either treated with OM-89, infected or treated and infected. For infection, monolayers were infected with TS4 at roughly 1*10^6^ CFU/ml (effective 1*10^5^ CFU/well/100 µl) for 4 hours. 1 hour into infection, FITC-labelled ampicillin at 64.5 µg/ml was added. Cells were washed several times with excess amounts of PBS washes before adding DMEM/F-12 with 20 mM HEPES omitting PBS or OM-89 and taking images on a Confocal Stellaris 5 using a HC PL APO 63x (NA 1.40) oil-immersion objective. To reduce low-frequency background variations, a Gaussian-blurred duplicate of the projection was subtracted from the original image. Background correction was performed using separate control images acquired under identical microscope settings in the absence of the labeled compound. Intracellular fluorescence was quantified after background subtraction of images of PBS or OM-89-treated cells using fiji script *mean_fluorescence*.

For antibiotic uptake per cell type, cells were treated as described above but instead of live-imaging, cells were fixed to stain for cell-specific cytokeratin markers. Briefly, cells were fixed in 4% PFA for 30 min at 4°C. After permeabilization, cells were incubated for 15 min in quenching buffer (2.5M glycine in PBS) before washing and blocking. Quenching was performed to reduce autofluorescence in the labelled antibiotic channel due to fixing. Cells were stained with primary antibodies against CK13 for intermediate cells and CK20 for umbrella-like cells. Primary antibodies were diluted 1:100 in 1% BSA in PBS. After washing, cells were incubated with 1:1000 dilution of secondary antibody, phalloidin-750 (Thermo Fisher) and DAPI in 1% BSA in PBS. Each staining was carried out for 1 hour at room temperature. All used antibodies are listed in Table 4. Images were taken on a Confocal Stellaris 5 using a HC PL APO 63x (NA 1.40) oil-immersion objective. To reduce low-frequency background variations, a Gaussian-blurred duplicate of the projection was subtracted from the original image. Binary masks were generated from the corresponding cell-type marker channel after thresholding on image to segment marker-positive cells using fiji script *cell mask_for_fluorescence*. The resulting masks were converted to binary images and used to define the cellular area of interest for cell-specific intracellular fluorescence quantification after multiplication with the binary cell masks using fiji script *mean_fluorescence_cell mask*. Background correction was performed using separate control images acquired under identical microscope settings in the absence of the labeled compound using fiji script *background_mean_fluorescence_cell mask*. For these controls, the same cell-type masks were applied, and mean fluorescence values were calculated. Cell-specific intracellular fluorescence was quantified from background-subtracted images of OM-89-treated cells.

#### Colocalization of intracellular UPEC with acidic compartments, Lamp1 and Cathepsin L

Differentiated mouse monolayers were infected with TS66 at roughly 1*10^6^ CFU/ml (effective 1*10^5^ CFU/well/100 µl) for 4 hours on a rocking platform to facilitate bacterial uptake for intracellular bacteria imaging. Thereafter, extracellular bacteria were washed with excessive amounts of PBS. For staining of acidic compartments, cells were incubated with LysoTracker DND-99 Red (Thermo Fisher) at 50 nM for 30 min following manufacturer’s recommendations. Thereafter, monolayers were fixed with 4% PFA for 1 hour at 4°C and stained with phalloidin and DAPI as mentioned above. For staining of Lamp1-positive vesicles, cells were fixed and stained with Lamp1 antibody, phalloidin and DAPI as mentioned above. For staining of enzymatically active compartments, Cathepsin L activity stained using Magic Red^TM^ Cathepsin L kit (bio-rad) following manufacturer’s recommendations. After 3 hours of infection, the Magic Red^TM^ substrate and NucBlue^TM^ Live ReadyProbes™ Reagent (Hoechst 33342, Thermo Fisher) was added on top for 60 min. After washing 3 times, cells were stained with CellMask^TM^ Plasma Membrane Stain in Deep Red (Thermo Fisher) for 10 minutes. After washing again, cells were briefly fixed with 4% PFA for 15 min at RT. Images were taken on a Confocal Stellaris 5 using a HC PL APO 63x (NA 1.40) oil-immersion objective. Intracellular bacteria were cropped by segmenting on bacterial signal inside the phalloidin signal of the bladder epithelial cells in fiji using the orthogonal view. Colocalization of intracellular bacteria with either LysoTracker, Lamp1 or Cathepsin L signal was then analyzed using the fiji script *fluorescent_overlap_intracellular_bacteria* by drawing a freehand region of interest (ROI) around bacteria and extracting the mean fluorescence intensity normalization over the area of the ROI. For random control ROIs, images were processed the same way except for moving the ROI from the bacterial cluster to an area without bacterial signal.

### Intraorganelle pH

The intraorganelle pH of differentiated monolayers was estimated using LysoSensor™ Yellow/Blue DND-160 (Thermo Fisher) following manufacturer’s recommendations with minor modifications. Briefly, after infection with TS4 as mentioned above, LysoSensor was diluted to a final concentration of 5 µM in live cell imaging solution (140 mM NaCl, 2.5 mM KCl, 1.8 mM CaCl_2_, 1.0 mM MgCl_2_, 20 mM HEPES, pH 7.4). Mouse and human bladder epithelial cells were incubated for 10 min and 20 min with the sensor, respectively, before measuring fluorescent intensity on a plate reader (Tecan Infinite M Nano) taking four measurements per well at Ex/Em of 329nm/440nm 348nm/540nm.

### Cathepsin assay

Cathepsin L activity in differentiated and TS4 infected monolayers was measured using Magic Red^TM^ Cathepsin L kit (bio-rad) following manufacturer’s recommendations. Cells were incubated with Magic Red^TM^ substrate for 60 min and fluorescent intensity was measured on a plate reader (Tecan Infinite M Nano) taking four measurements per well at Ex/Em of 592nm/628nm.

### RNA extraction, RT-qPCR and sequencing

#### Human bladder organoids

hBOs were extracted from the BME using Cell Recovery Solution. For RNA extraction, organoids were incubated with the appropriate volume of RNA lysis buffer (RNAeasy Plus Mini Kit, QIAGEN) and RNA was isolated following the manufacturer’s instructions. 250 ng of RNA were used to generate cDNA using the SuperScript®IV First-Strand Synthesis System with random hexamers (Invitrogen). RT-qPCR primer sequences for *UPK3A*, *KRT20*, *KRT13*, *TP63, ZO-1* and *Gapdh* are listed in Table 2. RT-qPCR reactions were prepared with SYBRGreen PCR Master Mix (Applied Biosystems) with 500 nM primers, and 1 µl cDNA. Reactions were run for quantification on QuantStudio 6 or 7 Flex Real-Time PCR System (Thermo Fisher) and amplicon specificity was confirmed by melting-curve analysis.

#### Mouse bladder organoids

mBOs were either left uninfected or were infected with TS85 as described above. Four hours after microinjection, the organoids were extracted from the BME using Cell Recovery Solution. For RNA extraction, organoids were incubated with the appropriate volume of RNA lysis buffer (RNAeasy Plus Mini Kit, QIAGEN) and RNA was isolated following the manufacturer’s instructions. RNA quantification and quality control was performed on a TapeStation 4200 (Agilent). Libraries were prepared using the “Illumina stranded mRNA ligation” (ISML) prep, according to Illumina protocol 1000000124518 v01. To account for bacterial mRNA contaminations during the mRNA polyA capture step, 200 ng of RNA for the PBSinf samples was used due to the presence of a double set of rRNA peaks whereas for all other samples 125 ng of RNA was taken. Libraries were then sequenced on the NovaSeq 6000 (Illumina) yielding approximately 25-35 million, 60-bp, paired end reads for all samples.

### RNA sequencing analysis

STAR (v.2.7.10b) (Dobin et al, 2013) was used to align FASTQ reads to the mouse mm10 reference genome and count reads in genes from the annotations in Ensembl release 102. Genes were considered expressed and maintained for downstream analysis if they had more than 10 counts in at least 4 of the samples (14,114 genes remaining). Differential expression analysis was performed using DESeq2 (version 1.36.0) and genes were considered differentially expressed if they were changed by more than 2-fold (log2FC ≥ 1) and had a corrected p-value ≤ 0.05. Two methods were used for exploring affected pathways: first, an over-representation analysis, using the g:Profiler (Kolberg et al, 2023) R client, was used to determine which pathways were enriched for up and down-regulated genes separately. For the gene set enrichment analysis, all expressed genes were ranked by their wald statistic and analyzed using the fgsea toolset in R (Korotkevich et al, 2016) for both GO and KEGG databases.

### Genebridge MMAS analysis

Genebridge MMAS (Module-Module Association Score) within the GeneBridge platform was used to identify and quantify connections between different biological modules (Li et al, 2019). Briefly, GeneBridge uses cross-species transcriptome compendia and statistical analysis to determine the significance of relationships between modules. We analyzed the KEGG lysosomal category to identify models positively correlating with it in eight human bladder datasets (dataset identifier: GSE13507 – sample size 265, GSE31189 – sample size 92, GSE48276 – sample size 84, GSE32894 – sample size 308, GSE83586 – sample size 307, GSE86411 – sample size 132, GSE32548 – sample size 131, GSE31684 – sample size 93).

### Statistical analysis

Statistics were performed with GraphPad Prism 10.3.1. Data and corresponding test results (e.g., normality results, T-values, degrees of freedoms, p-values) can be found in Supplementary Information.

## Data availability

Raw RNA-sequencing data have been deposited in the Gene Expression Omnibus (GEO) under accession number GSE306613. All other data supporting the findings of this study, including processed datasets and linked statistical analyses, will be made available on Zenodo (DOI: 10.5281/zenodo.17413104).

## Acknowledgements

We thank Dr. Ophélie Rutschmann and Mathilde Morelli for experimental and technical support; Dr. Naomi Wieser and Dr. Edouard Baulier from OM Pharma for helpful discussions and valuable feedback on the manuscript, as well as all of the McKinney laboratory for scientific discussions; the Gene Expression Core Facility at EPFL for RNAseq data generation; and the BioInformatics Competence Center, University of Lausanne and EPFL for analysis of the RNAseq data. Pre-submission review was conducted using qed Science (https://www.qedscience.com). OM Pharma providing OM-89.

## Funding

Behörden der Schweizerischen Eidgenossenschaft – Swiss Government Excellence Scholarship 2022.0289 (KT)

Swiss National Science Foundation (SNSF) – Postdoctoral Fellowship TMPFP3_217144 (KT)

IBSA Foundation – IBSA Foundation Fellowships Call 2021 in Fertility/Urology (KT) Fondation Leenaards (JM)

Swiss National Science Foundation (SNSF) – National Centre of Competence in Research (NCCR) AntiResist grant 51NF40_180541 (JM)

OM Pharma (JM)

## Competing interests

MR was and CP is an employee of OM Pharma SA, Meyrin, Switzerland. All other authors declare that they have no conflict of interest.

## Authors contribution

Conceptualization: KT, CP, MR, JDM

Data curation: KT, KS, GP, LS, VB, AMB

Formal analysis: KT, KS, GP, VB, AMB

Funding acquisition: KT, CP, JDM

Investigation: KT, KS, GP, LS, VB, AMB

Methodology: KT, MR

Project administration: KT, MR

Resources: CP, JDM

Supervision: KT, MR, CP, JDM

Validation: KT

Visualization: KT, AMB, MR

Writing original draft: KT

Writing review & editing: KT, KS, GP, AMB, VB, CP, MR

## Declaration of generative AI and AI-assisted technologies in the manuscript preparation process

During the preparation of this work the author(s) used ChatGPT (OpenAI) in order to assist with language refinement and clarity in sections of the manuscript, including improving phrasing, grammar, and flow. After using this tool/service, the author(s) reviewed and edited the content as needed and take(s) full responsibility for the content of the published article.

---

**SI Figure 1.**
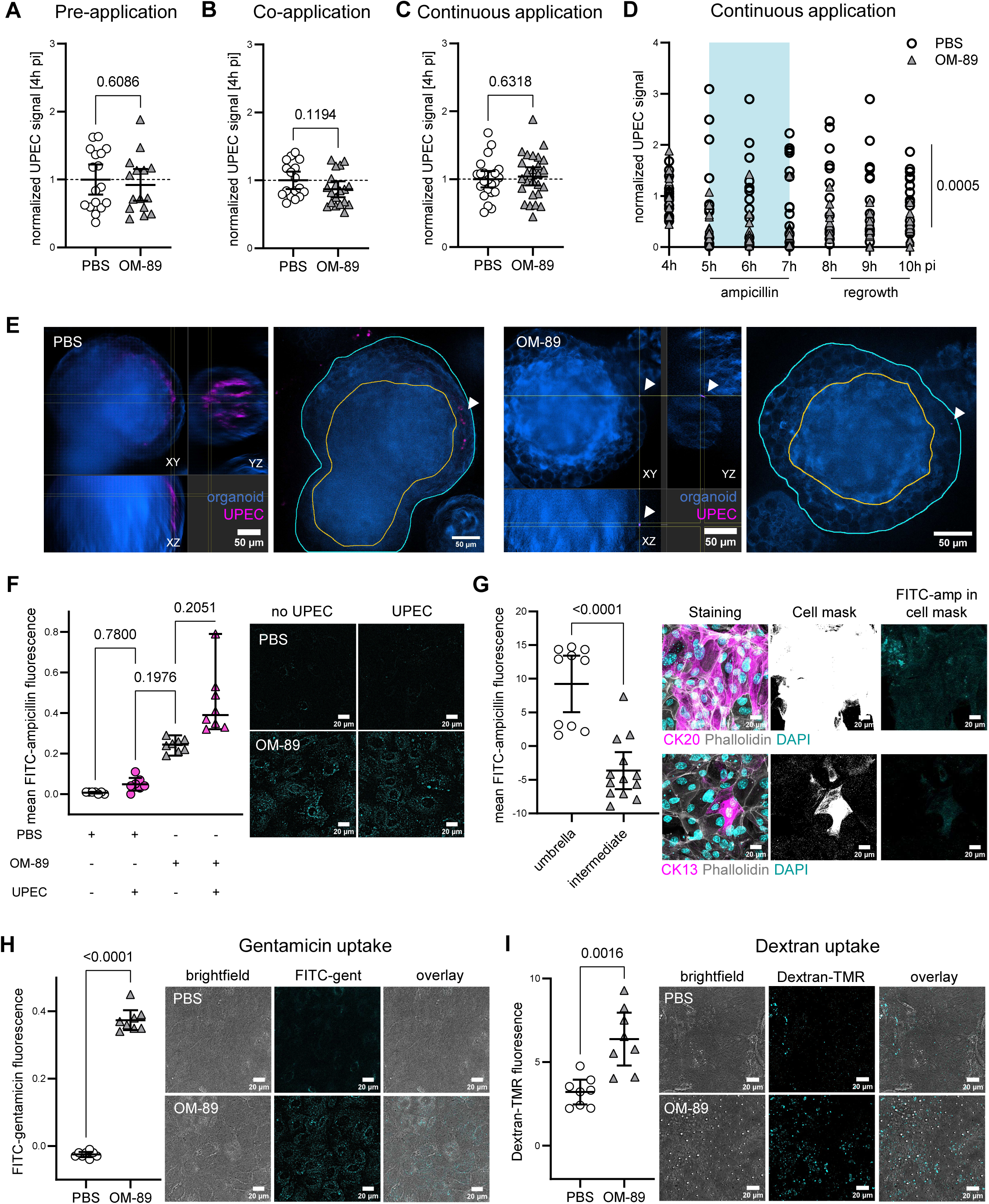
OM-89 enhances antibiotic uptake in bladder epithelial cells. (A-C) Mouse bladder organoids treated with OM-89 under (A) pre-application, (B) coapplication, or (C) continuous application regimes. Quantification of CFT073 fluorescence area at 4h post-infection (pi). Each dot represents one organoid. Values normalized to PBS control (dashed line). Mean ± 95% CI. Welch’s t test. N ≥ 15 per condition for (A), n ≥ 17 per condition for (B) and n ≥ 23 per condition for (C). (D) CFT073 signal over time in the continuous application regime. Quantification of bacterial fluorescence area normalized to PBS-treated organoids at the same timepoint. Each dot represents one organoid, lines connect matched organoids. RM two-way ANOVA with Šídák’s correction; p-value shown for treatment effect. N ≥ 14 per condition. (E) Cross-sectional (left) and single central plane (right) images of organoids with regrowing UPEC from the tissue 3h after antibiotic removal (10h pi) in the continuous treatment regime. Organoids shown in blue, UPEC shown in magenta. For central plane images: organoid boundaries indicated in cyan, luminal boundaries indicated in yellow. White arrow heads indicate UPEC regrowing in the organoid wall. (F) Uptake of FITC-labelled ampicillin in mouse bladder epithelial cell monolayers with or without UPEC infection. Quantification of fluorescence after background subtraction of PBS- or OM-89-treated cells according to infection status without labeled antibiotic. Mean ± 95% CI. Kruskal-Wallis test with Dunn’s correction. N ≥ 6 per condition. Z-projection (maximum intensity) of representative images. FITC-ampicillin in cyan. For uninfected cells, values and images correspond to Figure 1 I. (G) Uptake of FITC-labelled ampicillin in OM-89 treated mouse bladder epithelial cell monolayers according to cell type. For each cell type, a binary cell mask was generated from the corresponding cell-type marker channel (CK20+ for umbrella cells, CK13+ for intermediate cells) and used to restrict measurements to the cellular area. Quantification of fluorescence after background subtraction of images without labelled antibiotics where the same cell-type masks were applied. Mean ± 95% CI. Mann-Whitney test. N ≥ 10 per condition. Z-projection (maximum intensity) of representative images. CK20 and CK13-cell staining in magenta, phalloidin in grey, DAPI in cyan. Cell mask generated from the cellspecific stainings in white. Masked areas of FITC-ampicillin in cyan. (H), (I) Uptake of (H) FITC-labeled gentamicin and (I) Dextran-TMR in mouse bladder epithelial cell monolayers. Quantification of fluorescence after background subtraction of PBS- or OM-89-treated cells without labeled antibiotic. Mean ± 95% CI. Welch’s test. N = 8 per condition. Z-projection (maximum intensity) of representative images. FITC-gentamicin and Dextran-TMR shown in cyan.

**SI Figure 2.**
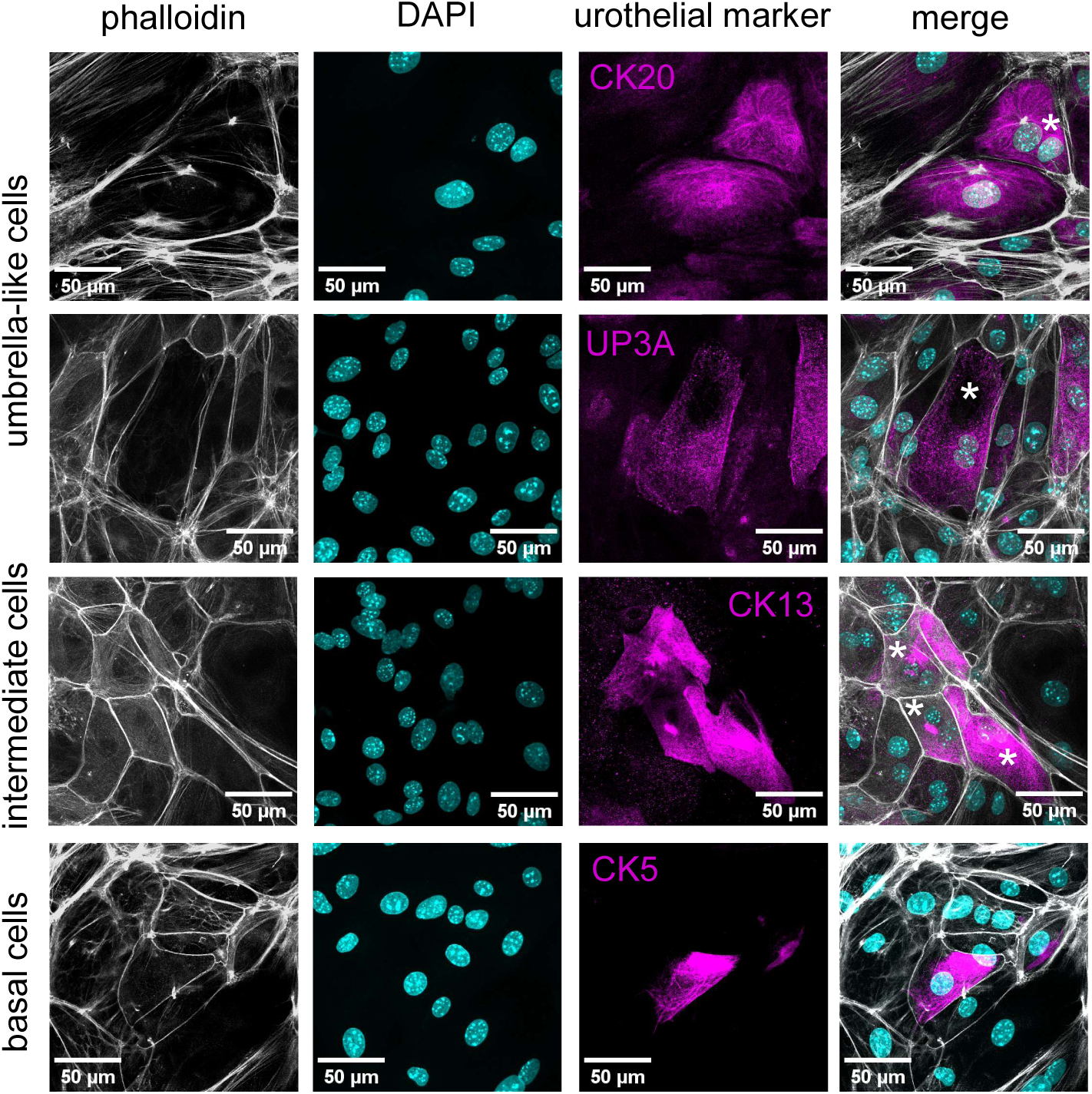
Establishment of differentiated mouse bladder epithelial cell monolayer. Staining of differentiated mouse monolayer for bladder epithelial cell markers: Cytokeratin (CK)20 – umbrella-like cells. Uroplakin (UP)3A – umbrella-like cells. CK13 – intermediate cells. CK13-positive, binucleated cells indicate late intermediate cells. CK5 – basal cells. Binucleated cells indicated with asterisks.

**SI Figure 3.**
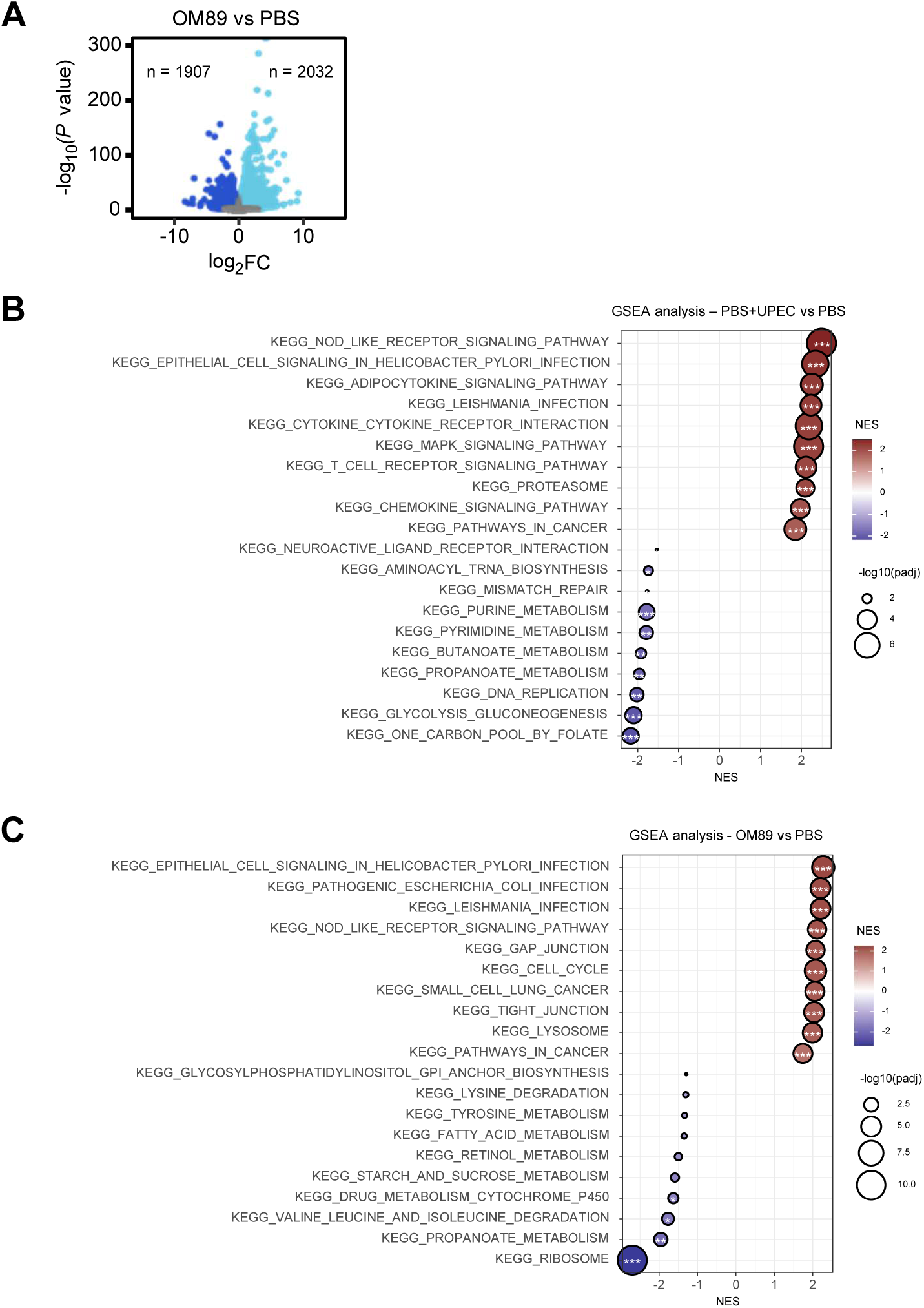
Transcriptomic analysis of OM-89-treated bladder organoids. (A) Volcano plots of differentially expressed genes (log2FC ≥ 1; adjusted p ≤ 0.05) comparing OM-89- (cyan) and PBS-treated organoids without infection (dark blue). (B) Gene set enrichment analysis (GSEA) of the top 10 Kyoto Encyclopedia of Genes and Genomes (KEGG) terms comparing infection vs no infection in PBS control organoids. NES, normalized enrichment score. Ns changes not highlighted. (C) GSEA of the top 10 KEGG terms comparing OM-89- and PBS-treated organoids without infection. Ns changes not highlighted (A-C) Data from four independent wells with n ≥ 50 organoids each.

**SI Figure 4.**
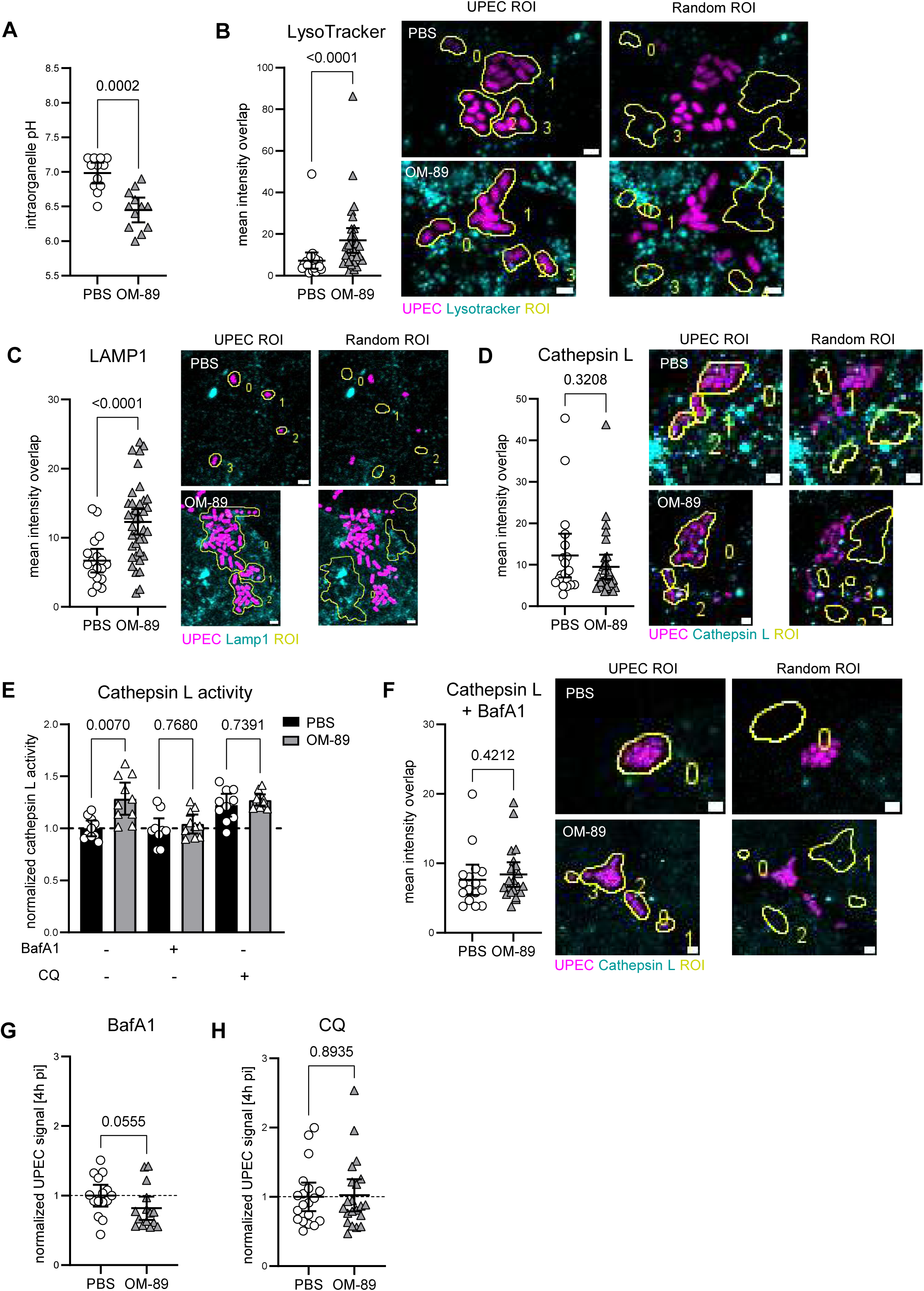
Controls and complementary analyses for lysosomal activity and blocking experiments. (A) Intraorganelle pH in uninfected monolayers with and without OM-89 treatment. Mean ± 95% CI. Mann-Whitney test. N = 12 per condition. (B)-(D), (F) Quantification of colocalization of random regions of interest (ROIs) with (B) LysoTracker, (C) LAMP1, (D) Cathepsin L and (F) Cathepsin L with bafilomycin A1 (BafA1) in monolayers of mouse bladder epithelial cells. Intensities normalized to random ROI area. (B), (D), (F) Mann-Whitney test. (C) Welch’s test. N ≥ 24 per condition for (B), n ≥ 18 per condition for (C), n ≥ 19 per condition for (D) and n ≥ 16 per condition for (F). Zprojection (maximum intensity) of inserts in XYZ cross-section images shown in Figure 4D-G. UPEC, magenta; LysoTracker/LAMP1/Cathepsin L, cyan. ROIs drawn around intracellular UPEC used for intensity quantification of specific/active co-localization in Figure 4D-G. Random ROIs used for unspecific/random co-localization. Scale bar 2 μm. (E) Cathepsin L activity during infection with or without OM-89, and with BafA1 or chloroquine (CQ) blocking. Values normalized to unblocked PBS controls (dashed line). Mean ± 95% CI. Brown-Forsythe ANOVA with Welch’s correction. N = 10 per condition. (G), (H) CFT073 growth after (G) BafA1 and (H) CQ blocking. Quantification of bacterial fluorescence area inside organoids 4h pi. Each dot represents one organoid. Values normalized to PBS control (dashed line). Mean ± 95% CI. Mann-Whitney test. N ≥ 15 per condition for (G) and n ≥ 19 per condition for (H).

**SI Figure 5.**
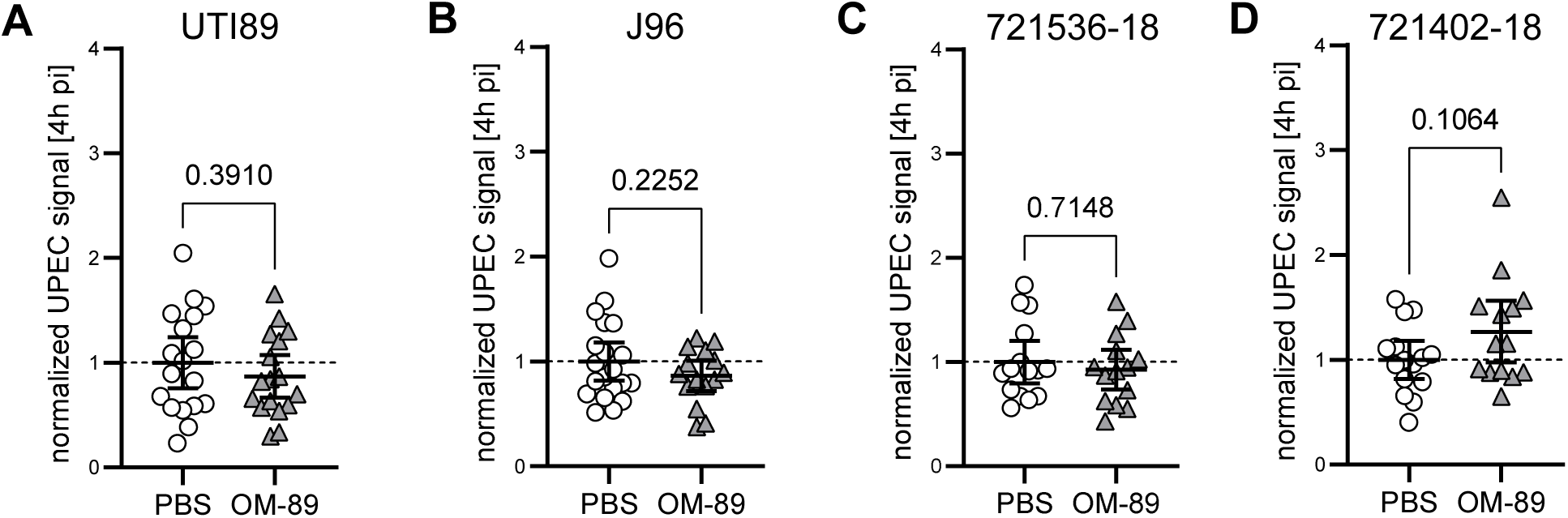
Growth of different UPEC strains in mouse bladder organoids. (A)-(D) Growth of UPEC strain (A) UTI89, (B) J96, (C) 721536-18 and (D) 721402-18. Quantification of bacterial fluorescence area inside organoids at 4h pi. Each dot represents one organoid. Values normalized to PBS control (dashed line). Mean ± 95% CI. Welch’s test for UTI89 (A), J96 (B), 721402-18 (D). Mann-Whitney test for 721536-18 (C). N ≥ 17 per condition for (A), n ≥ 15 per condition for (B), n ≥ 14 per condition for (C), (D).

**SI Figure 6.**
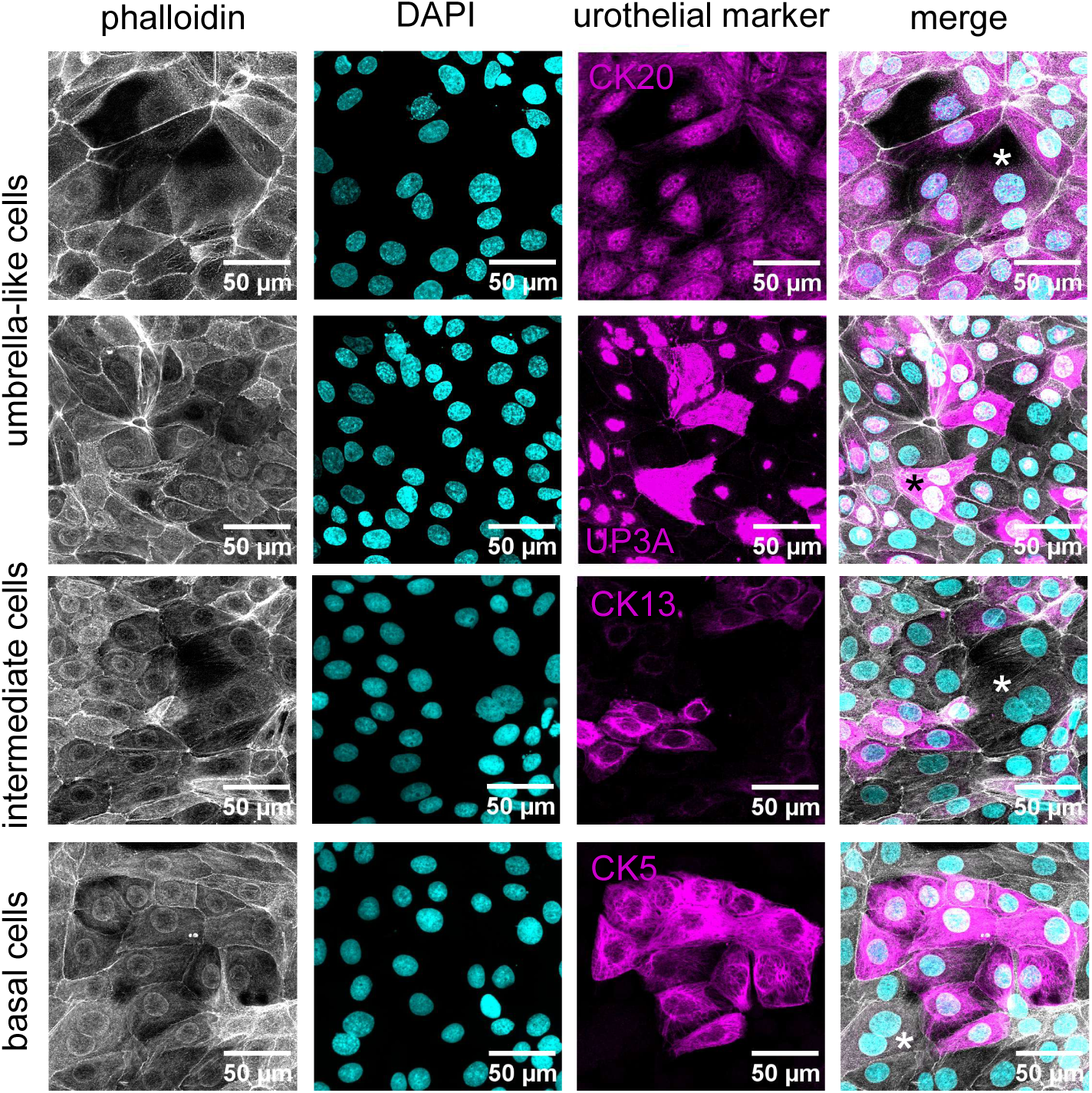
Establishment of differentiated human bladder epithelial cell monolayer. Staining of differentiated mouse monolayer for bladder epithelial cell markers: Cytokeratin (CK)20 – umbrella-like cells. Uroplakin (UP)3A – umbrella-like cells. CK13 – intermediate cells. CK5 – basal cells. Binucleated cells indicated with asterisks.

**SI Figure 7.**
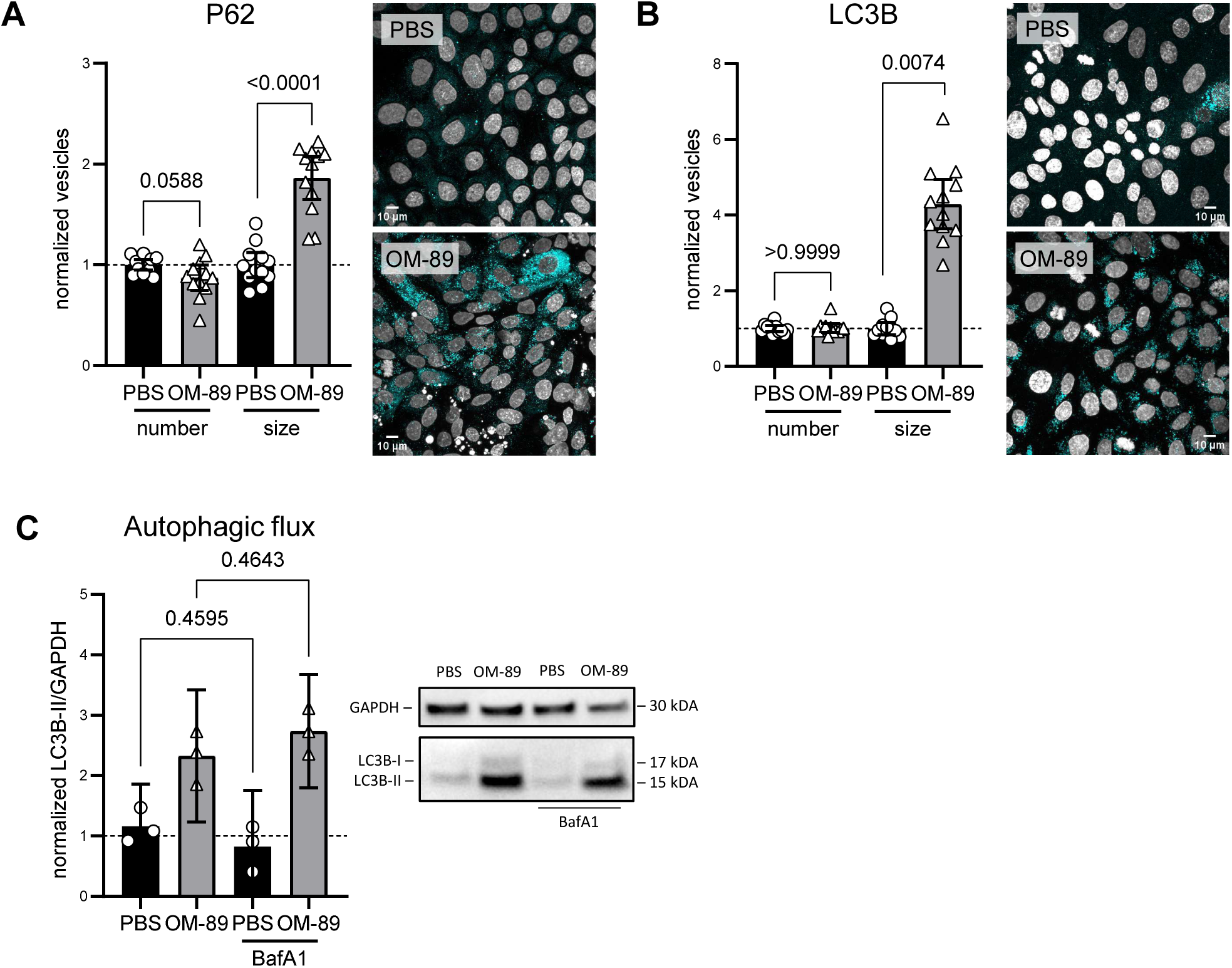
Autophagy-related markers and flux in human bladder epithelial monolayers treated with OM-89. (A), (B) Quantification of vesicle numbers and size for (A) P62 and (B) LC3B in infected monolayers of human bladder epithelial cells with and without OM-89 treatment. Values normalized to PBS control (dashed line). Mean ± 95% CI. Welch’s t test for P62 vesicle counts (A) and LC3B vesicle size (B). Mann-Whitney test for P62 vesicle size (A) and LC3B vesicle counts (B). N ≥ 11 per condition. Z-projection (maximum intensity) of representative images. Foci, cyan; DAPI, grey. (C) Western blot analysis of LC3B during infection of human monolayers. LC3B-II/GAPDH ratio normalized to PBS control (dashed line). Mean ± 95% CI. Brown-Forsythe ANOVA with Welch’s correction. N = 3 per condition.

**SI Figure 8.**
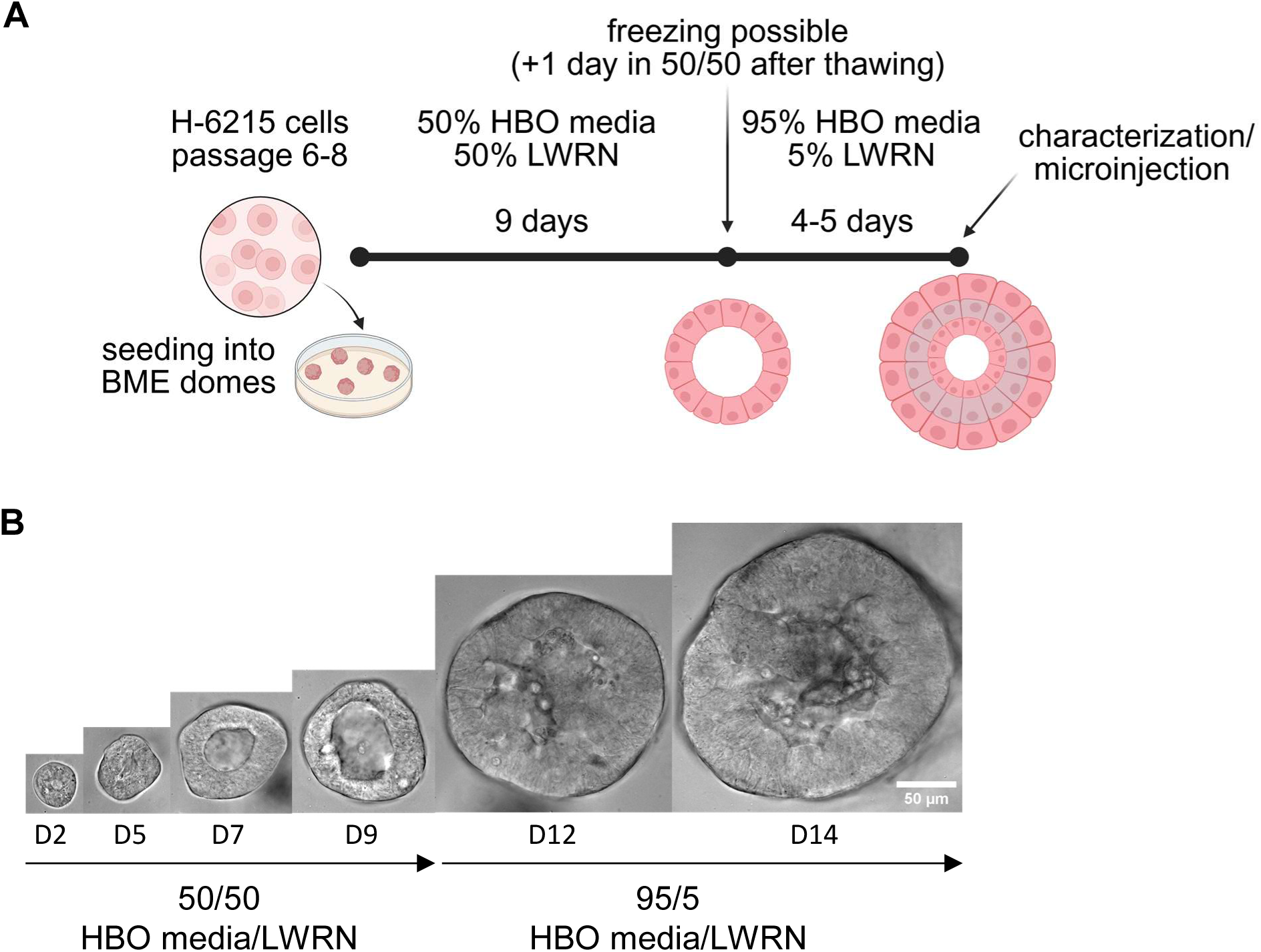
Establishment of human bladder organoids from primary human bladder epithelial cells. (A) Schematic of the human bladder organoid culture protocol. Primary H-6215 bladder epithelial cells (passages 6-8) were seeded as single cells into basement membrane extract (BME) domes and cultured in medium containing 50% HBO medium and 50% LWRN-conditioned medium (50/50). After 9 days, organoids could be cryopreserved. Following thawing, organoids were recovered for 1 day in 50/50 HBO/LWRN medium before switching to differentiation conditions (95% HBO medium with 5% LWRN, 95/5). Differentiated human bladder organoids were characterized or used for microinjection experiments on days 13-14. Created with BioRender.com. (B) Growth of human bladder organoids over time. H-6215 cells embedded in BME rapidly expand and form organoids with a central lumen visible as early as 2 days after seeding. Images show representative examples from different organoids.

**SI Figure 9.**
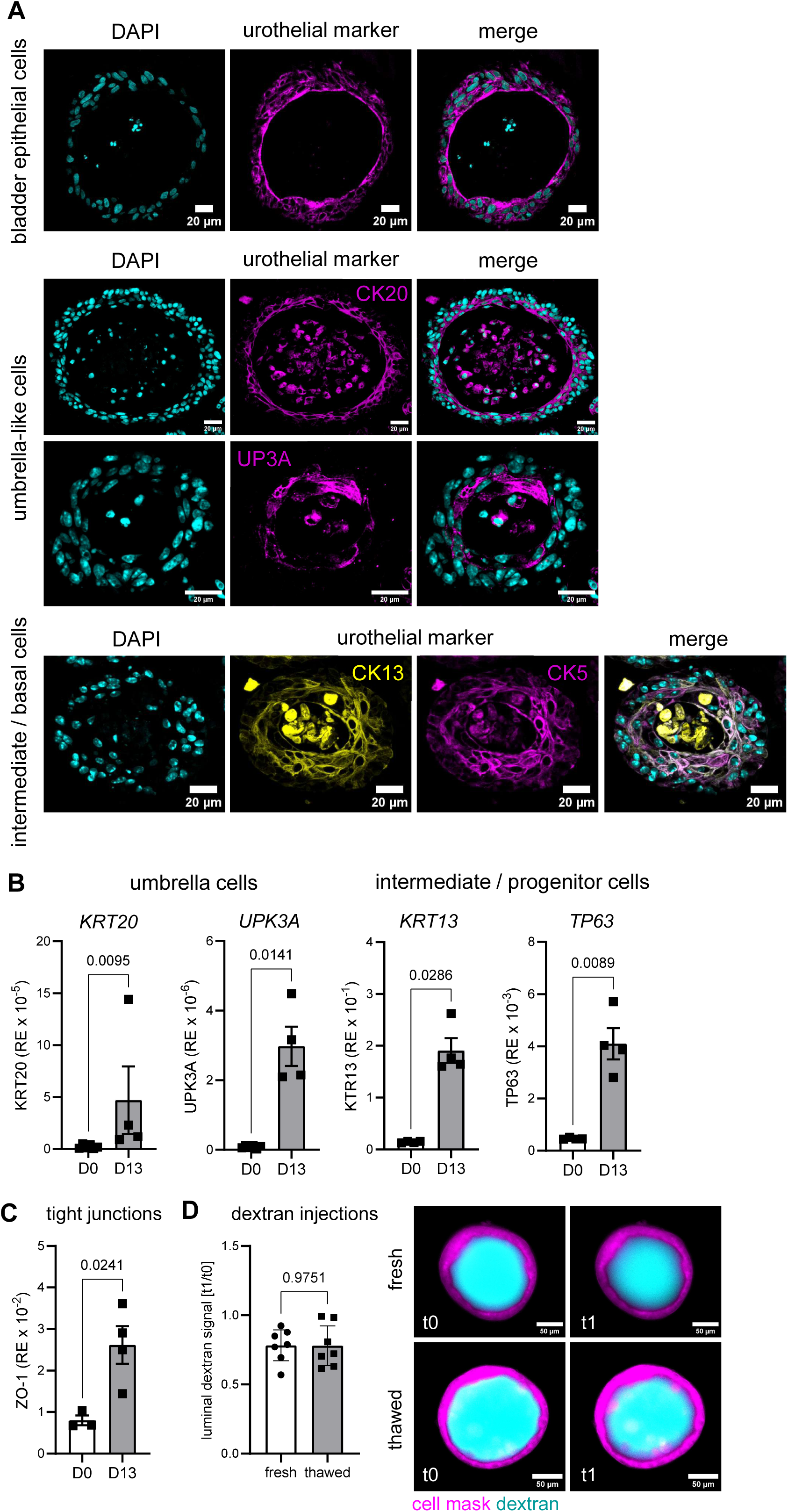
Human bladder organoids express urothelial markers and form a tight lumen. (A) Immunostaining of stratified and differentiated human bladder organoids derived from primary H-6215 cells for urothelial markers. Cytokeratin (CK)7 marks general urothelium; CK20 and uroplakin (UP)3A mark umbrella-like cells; CK13 marks intermediate cells; CK5 marks basal cells. Representative images of paraffin-embedded sections are shown. For CK20 and UP3A, two organoids are displayed. Organoids were fixed for staining on day 13 of differentiation. (B) RT-qPCR analysis of differentiation markers in human bladder organoids. Differentiated organoids (day 13, D13) upregulate umbrella cell markers (KRT20, UPK3A), intermediate cell markers (KRT3), and progenitor cell markers (TP63) compared to undifferentiated H-6215 monolayers (day 0, D0). Mann-Whitney test for KRT20 and KRT3. Welch’s t test for UPK3A and TP63. N ≥ 4. (C) RT-qPCR analysis of tight junction protein ZO-1 in human bladder organoids. Differentiated organoids (day 13, D13) upregulate ZO-1 compared to undifferentiated H- 6215 monolayers (day 0, D0). Welch’s t test. N ≥ 3. (D) Dextran microinjection assay confirming luminal barrier integrity in fresh and thawed organoids. Dextran fluorescence was quantified in the central plane at t0 (approximately 30 min post-injection) and t1 (t0 + 1h). Organoids were injected either after continuous differentiation from single cells (fresh) or after freezing at day 9 followed by thawing and differentiation (thawed). Each dot represents one organoid. Values normalized to t0 for each organoid. Mean ± 95% CI. Fold change in dextran signal from t0 to t1: fresh organoids, 0.78 ± 0.12; thawed organoids, 0.78 ± 0.16. N = 7. Representative images of central plane of fresh and thawed organoids. Cell mask, magenta; Dextran, cyan.

**SI Figure 10.**
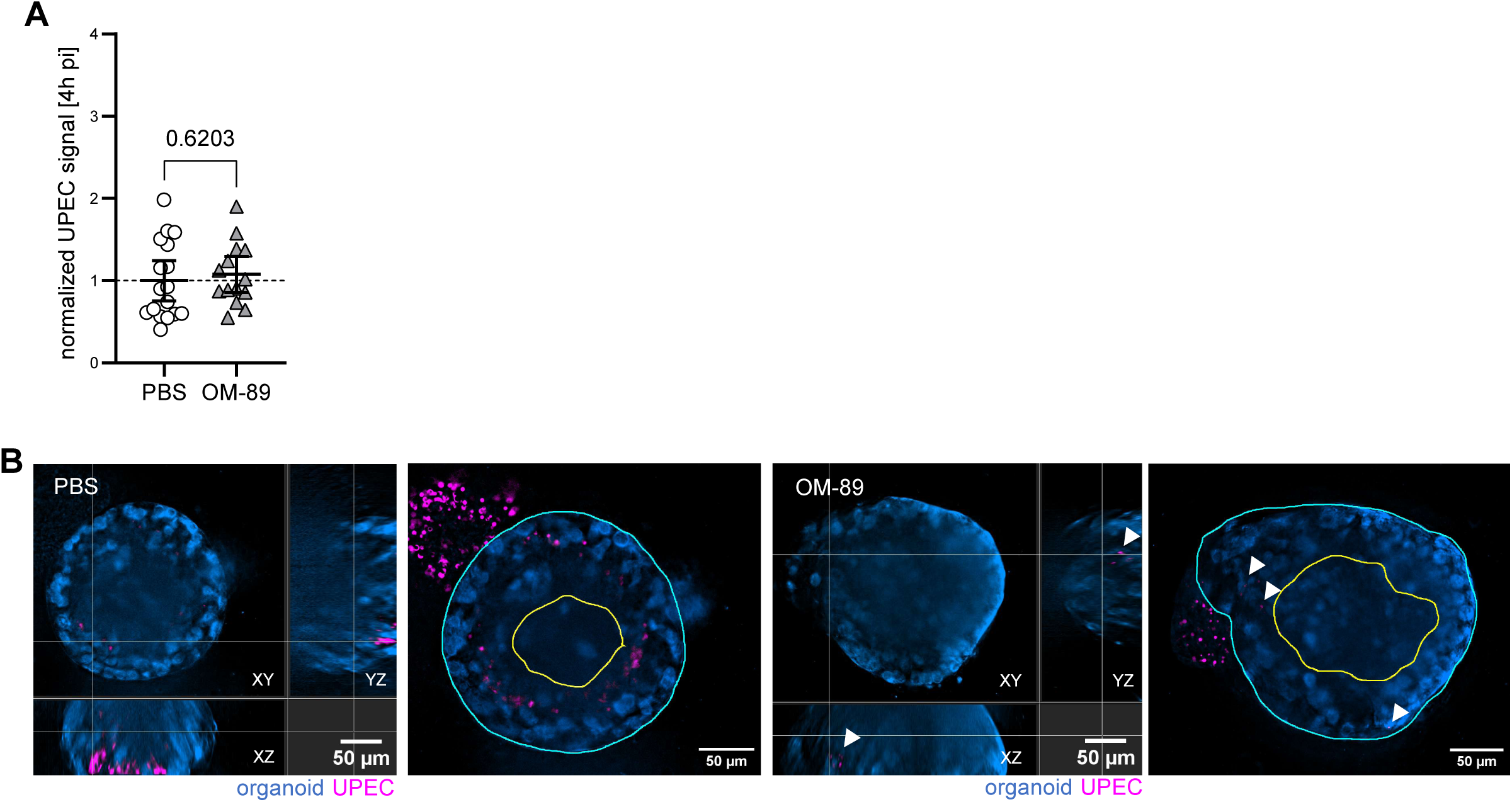
Growth and regrowth of UPEC in human bladder organoids. (A) Growth of UPEC strain CFT073. Quantification of bacterial fluorescence area inside organoids at 4h pi. Each dot represents one organoid. Values normalized to PBS control (dashed line). Mean ± 95% CI. Welch’s test. N ≥ 14 per condition. (B) Cross-sectional (left) and single central plane (right) images of human organoids with regrowing UPEC from the tissue 3h after antibiotic removal (10h pi) in the continuous treatment regime. Organoids shown in blue, UPEC shown in magenta. For central plane images: organoid boundaries indicated in cyan, luminal boundaries indicated in yellow. White arrow heads indicate UPEC regrowing in the organoid wall.

## Notes

### Competing Interest Statement

M.R. was and C.P. is an employee of OM Pharma SA, Meyrin, Switzerland. All other authors declare that they have no conflict of interest.

### Summary of Updates

This version of the manuscript has been revised to improve clarity, strengthen the mechanistic framework and better contextualize the translational relevance of the study. The Introduction and Discussion were updated to more clearly position OM-89 as a clinically approved therapy with long-standing use and previously unresolved mechanisms of action. Text has been refined throughout to improve readability and conceptual flow. The Results section has been revised for clarity and consistency, including improved description of experimental design. Additional data, especially on human epithelial models and primary clinical isolates, have been incorporated. Figure annotations and legends have been updated accordingly. Minor corrections have been made throughout the manuscript, including grammar, terminology consistency and formatting. Author affiliations and supplementary materials have also been updated.

